# AMOEBA Polarizable Molecular Dynamics Simulations of Guanine Quadruplexes: from the c-Kit Proto-oncogene to HIV-1

**DOI:** 10.1101/2024.08.28.610081

**Authors:** Dina El Ahdab, Louis Lagardère, Zeina Hobaika, Théo Jaffrelot Inizan, Frédéric Célerse, Nohad Gresh, Richard G. Maroun, Jean-Philip Piquemal

## Abstract

Long oligomer sequences, rich in guanine and cytosine, such as *c-kit1* and the HIV-1 LTR-III sequence, are prevalent in oncogenes and retroviruses and play crucial roles in cancer. Understanding the conformational dynamics of such guanine quadruplexes and identifying druggable regions are therefore essential for developing new inhibition strategies. In this study, we used extensive AMOEBA polarizable force field molecular dynamics simulations combined with data-driven adaptive sampling and clustering algorithms, reaching a cumulative simulation time of 7.5 *µ*s for *c-kit1*. Such simulations identified novel structural motives and show-cased the flexible loop dynamics, as well as the role of polarizable water in transient stabilization of the G-quadruplex. They also identified two druggable pockets in *c-kit1*. The 400 ns simulation of the HIV-1 LTR-III sequence confirmed its quadruplex stability and uncovered a potentially druggable cryptic pocket.

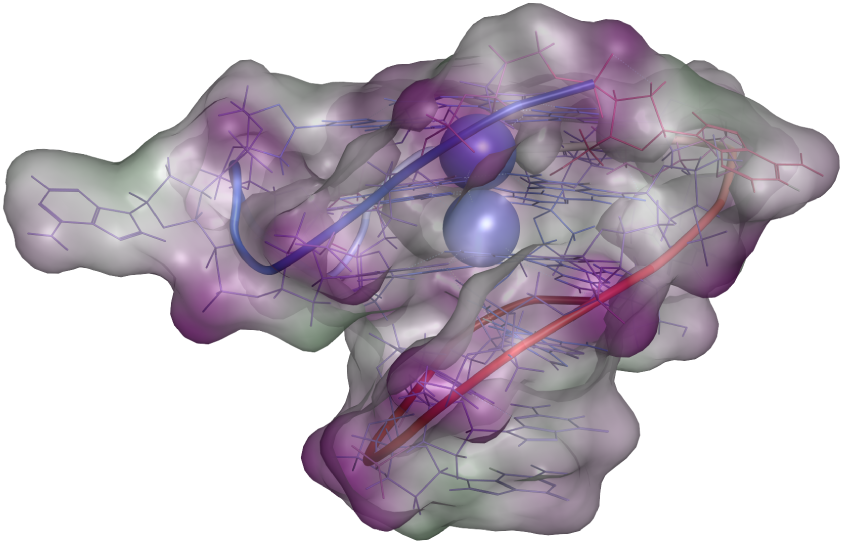

## Introduction

Long oligomer guanine-rich sequences are prevalent in oncogenes and retroviruses. ^1,2^ The tendency of these sequences to undergo a transition from double to quadruple helices has long been established.^3,4^ This transition occurs following strand separation and involves the formation of stacked guanine tetrads within the guanine-rich strand. ^5,6^ These tetrads are stabilized by Hoogsteen base pairing. In this arrangement, each guanine base utilizes its O6 and N7 atoms to accept protons from the N1 and N2 atoms, respectively, of an adjacent guanine, and reciprocally donates protons from its N1 and N2 atoms to the O6 and N7 atoms of another guanine.^6^ A common structural motif of guanine quadruplexes (GQs) consists of three stacked tetrads connected by one or more nucleotide loops, as illustrated in Figure 1. Alkali metal cations, primarily K^+^ and Na^+^, are essential for GQ stability.^7–10^ Typically, two cations are located along the axis perpendicular to the stacked tetrads, near their geometric centers.^11,12^ Several G-quadruplexes are recognized targets for chemotherapeutic ligand design, especially in the case of human telomeres.^2^ In cancer cells, the prevention of telomere shortening by telomerase is a key factor in cellular immortalization.^1^ Disrupting telomeric G-quadruplexes with ligands presents a strategy to induce telomere shortening and limit cancer cell proliferation.^13^ Therefore, inhibition of GQs by high-affinity ligands may impede immortalization.^1^ GQs participate in the regulation of gene expression at both transcriptional and translational levels.^1^ Furthermore, they are found in the promoter regions of various genes critical for cell signaling, and their overexpression may contribute to cancer development.^14–17^ In this study, we present multi-µs polarizable molecular dynamics simulations of two GQs, one human and one retroviral, to elucidate their distinct structural characteristics. The human GQ investigated is c-Kit, a proto-oncogene encoding the stem cell factor receptor CD117. CD117, a tyrosine kinase receptor, is involved in intracellular signaling pathways affecting differentiation, survival, adhesion, motility, angiogenesis, and proliferation.^18^ Mutations in CD117 can lead to several cancers, including gastrointestinal stromal tumor (GIST), acute myeloid leukemia, and melanoma.^19^ Although imatinib mesylate, a tyrosine kinase inhibitor, has demonstrated efficacy in treating GIST patients, inhibitor resistance has limited its optimal response.^18,20^ Therefore, targeting *c-kit* transcription to regulate CD117 mRNA expression represents a potential alternative therapeutic strategy.

**Figure 1:**
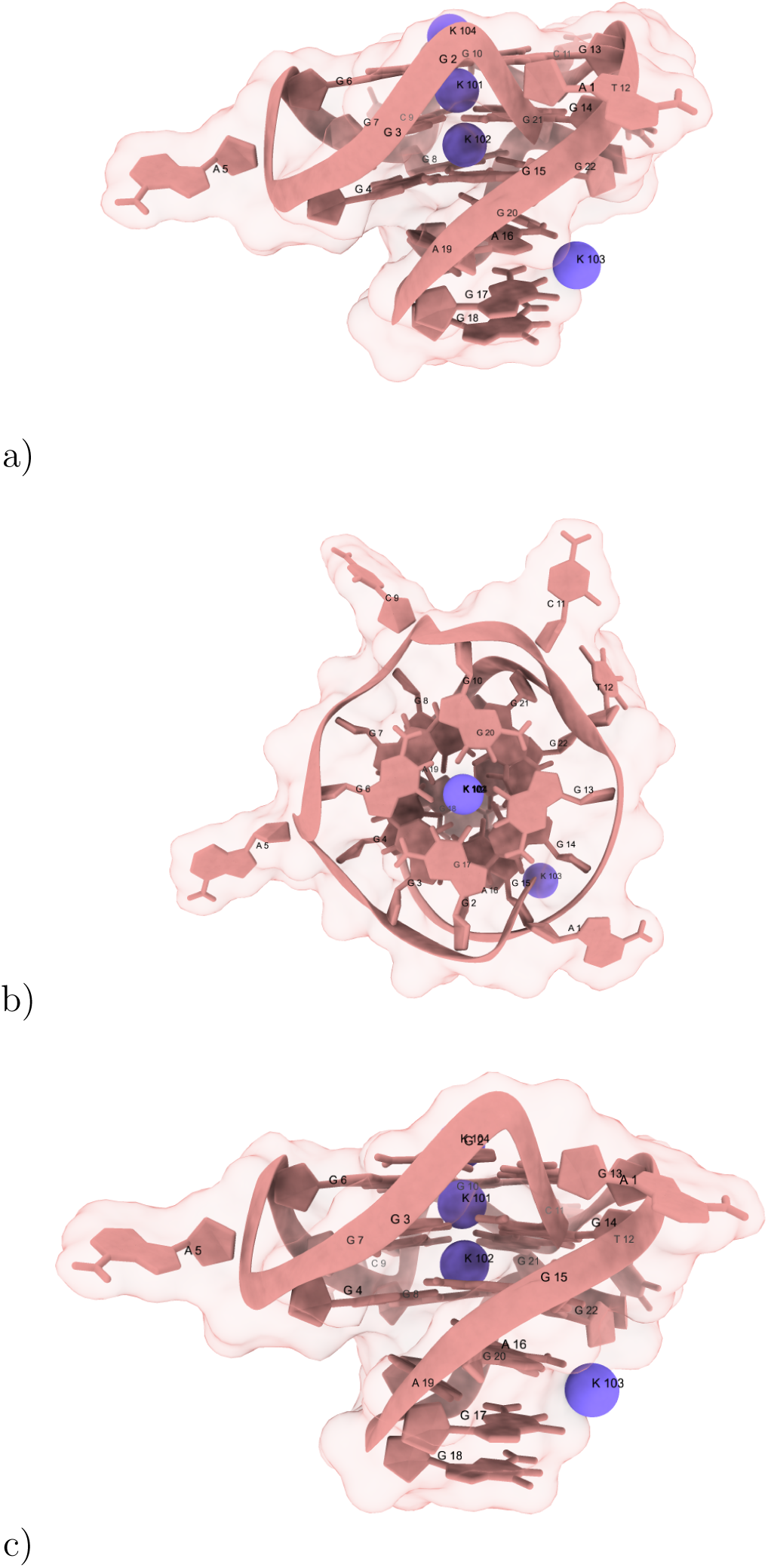
*c-kit1* GQ representation. *c-kit1* is formed by three G4 tetrad, one LP loop (A16, G17, G18, A19 and G20), two single nucleotide loops (A5 and C9), one two nucleotides loop (C11 and T12) and a flanking nucleotide (A1). a) General representation showing all the nucleotides of *c-kit1* GQ. b) A top view showing organization of G nucleotides around the ionic channel. c) View from the side for a better visibility of the LP loop.

Stabilizing the *c-kit1* proto-oncogene GQ with high-affinity ligands, by competing with polymerase binding, may represent a novel transcription-targeted therapeutic strategy against cancer. We focus on the *c-kit87up, 22-mer* sequence, d(AGGGAGGGCGCTGGGAGGAGGG), located 87 nucleotides upstream of the transcription start site of the *c-kit1* gene in the human genome. This sequence forms a single GQ species in K^+^-containing solutions.^21^ The presence of four guanine tracts, each consisting of three consecutive guanines and separated by linkers of one to four residues, results in the adoption of a canonical parallel-stranded GQ structure. As shown in Figure 1, the first tetrad is formed by guanine residues G2, G6, G10, and G13. The second tetrad is formed by G3, G7, G14, and G21. The third tetrad is formed by G4, G8, G15, and G22. NMR and circular dichroism experiments have demonstrated the essential role of the connecting linker sequences in GQ core stabilization.^21^ Four loop regions are present within the structure: single-nucleotide loops at A5 and C9, a dinucleotide lateral loop at C11 and T12, and a five-nucleotide loop from A16 to G20. This loop arrangement deviates from the typical three-propeller loop structure observed in parallel-stranded quadruplexes. Highresolution structures of the *c-kit1* GQ in the presence of K^+^ are available from 3D-NMR spectroscopy (PDB code 2O3M) and X-ray crystallography (PDB codes: 3QXR, 4WO2, and 4WO3).^11,22,23^

The crystal structure with the highest resolution (1.82 Å) for the 22-mer human native c-Kit proto-oncogene promoter, PDB code 4WO2, was selected as the starting structure for this study. This study is then extended to the G-quadruplexes of the HIV-1 long terminal repeat 3 (LTR-III). Two recent molecular dynamics (MD) simulations have investigated the LTR-III quadruplex using the non-polarizable AMBER 18 force field^24^ and the polarizable Drude 2017 force field. ^25^ The present study, employing the AMOEBA force field, will enable the elucidation of similarities and differences between the c-Kit and HIV-1 quadruplexes. Several guanine-rich sequences within the U3 promoter region of the HIV-1 long terminal repeat (LTR) have been shown to fold into a range of dynamically interconverting G-quadruplex structures.^26–28^ These sequences regulate viral transcription by stabilizing LTR G-quadruplexes. Among the G-quadruplexes formed within the LTR sequence, the LTR-III sequence exhibits the greatest conformational diversity in vitro. Consequently, an NMR structure of LTR-III in K^+^ solution revealed the following structural motifs: i) a unique quadruplex–duplex hybrid consisting of a three-layer (3 + 1) G-quadruplex scaffold with three strands (G15–G17, G19–G21, and G26–G28) oriented downward and one strand (G1–G2) oriented upward; ii) a 12-nucleotide diagonal loop containing a conserved duplex stem in which six nucleotides interact via Watson-Crick hydrogen bonds to form a hairpin; iii) a 3-nucleotide lateral loop; and iv) a 1-nucleotide propeller loop^29,30^ (Figure 2).

**Figure 2:**
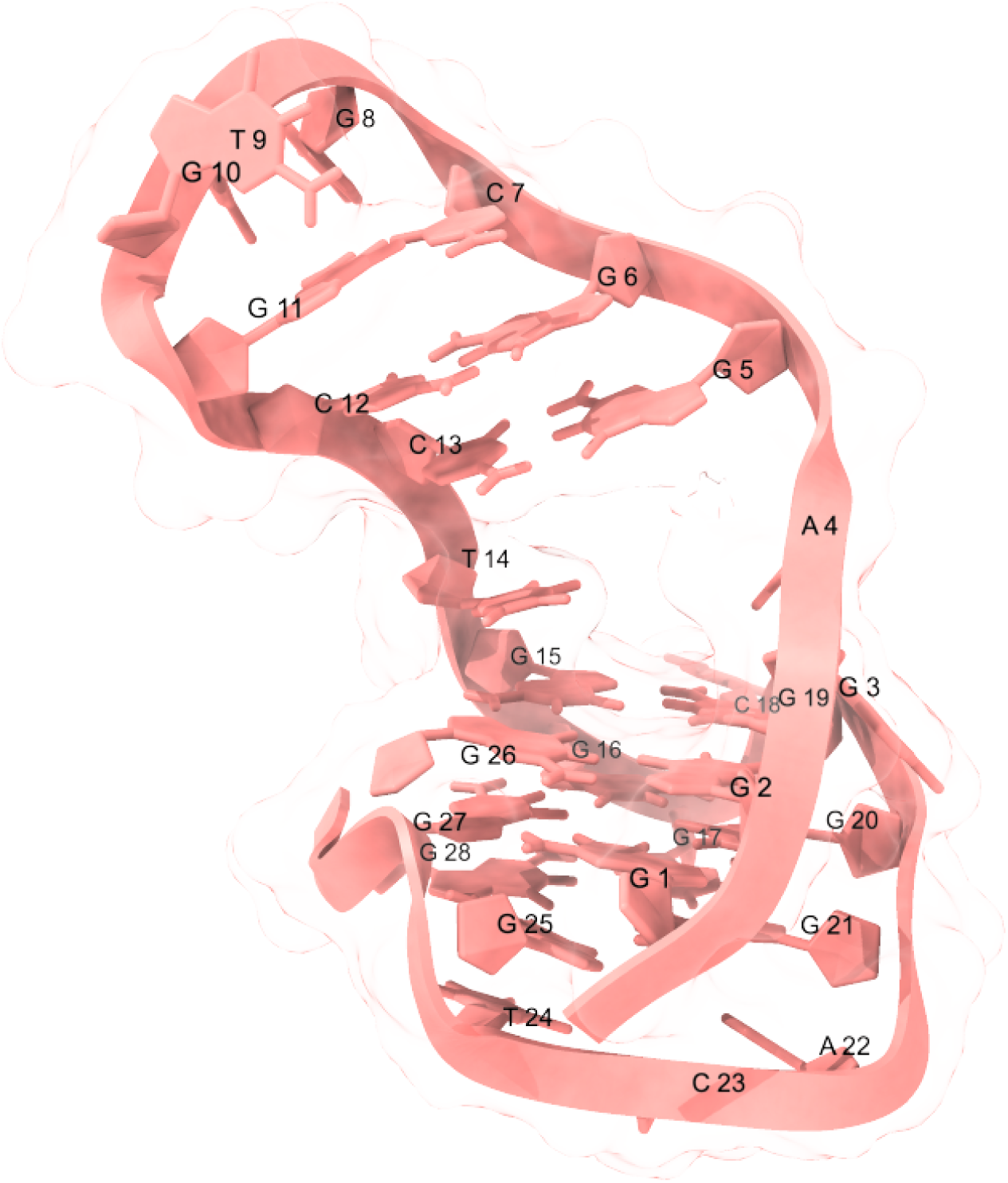
HIV-1 long-terminal repeat 3 (LTR-III) GQ representation showing the following motives: i) a unique quadruplex–duplex hybrid consisting of a three-layer (3 + 1) GQ scaffold with three strands (G15–G17, G19–G21, and G26–G28) pointing down and one strand (G1–G2) pointing up; ii) a 12-nt diagonal loop containing a conserved duplex-stem where six nucleotides are interacting by W-C hydrogen bonds to form a hairpin and iii) a 3-nt lateral loop and a 1-nt propeller loop.

X-ray crystallography and NMR spectroscopy provide high-resolution starting structures for subsequent molecular dynamics (MD) simulations. MD simulations, in turn, offer insights into the dynamics of the simulated macromolecule, the amplitudes of domain motions, and the potential occurrence of transient cryptic binding sites. Such information can be crucial for drug design. To this end, we employed the AMOEBA force field, specifically its extension for nucleic acids.^31^ This force field enables a reliable description of the stability conferred by the presence of two K^+^ cations bound within the central cavity of two successive stacked guanine tetrads. Indeed, polarizable force fields have been demonstrated to be particularly well-suited for simulating complex and highly charged biological environments.^32–38^ In this context, a recent validation study using the SIBFA polarizable potential^39^ is noteworthy. This study showed that SIBFA accurately reproduces the quantum mechanical (QM) energy profile for the translocation of alkali cations along the central axis of two stacked guanine tetrads,^40^ a phenomenon not readily captured by non-polarizable force fields.^41^

We employed the latest AMOEBA force field, implemented in the GPU-accelerated Tinker-HP software,^42,43^ in conjunction with unsupervised adaptive sampling. This strategy^35^ has been shown to significantly enhance the conformational sampling of the SARS-CoV-2 main protease^35,36^ and spike proteins,^38^ enabling access to multi-µs MD simulations. For GQs, this approach facilitates an in-depth understanding of the c-Kit G-quadruplex due to the extended simulation times achievable. Indeed, overstabilization of the quadruplex conformation of the*c-kit1* proto-oncogene could impede RNA polymerase-mediated initiation of transcription at an early genomic stage,^1^ potentially circumventing resistance to tyrosine kinase inhibitors, which occurs at the later proteomic stage.

AMOEBA simulations of c-Kit will be compared with recent MD results obtained using the AMBER12 force field^12^ and the additive CHARMM36 force field and its polarizable extension, Drude-2017.^30^ These comparisons reveal previously unobserved structural features of c-Kit, including differences in clustering, loop dynamics, cation responses within the ion channel, and the emergence of alternative transient states.

We will first investigate the c-Kit conformational landscape, including possible stable substates and the dynamics of loop rearrangements. Cluster analyses will be presented in this context. The extent of quadruplex stabilization by K^+^ and polarizable water molecules will then be analyzed. MD simulations will also be presented for an HIV-1 quadruplex, specifically the one within its LTR-III motif. We will conclude by elucidating potential cryptic pockets in both c-Kit and LTR-III.

## Results and discussion

AMOEBA molecular dynamics simulations of the *c-kit* G-quadruplex revealed a complex conformational landscape, characterized by multiple stable states and dynamic loop rearrangements. Cluster analysis identified distinct conformational states. The stabilizing effects of K^+^ ions and polarizable water molecules were examined. Simulations were extended to the LTR-III G-quadruplex, and a comparative analysis highlighted similarities and differences in the conformational dynamics and potential ligand binding sites of the two systems.

### Conformational stability and transition states

#### Evolution of the stability of the different GQ *c-kit1* nucleotides

Root mean square deviation (RMSD) was calculated for all of the 7.5 *µ*s of adaptive sampling trajectories to measure the average distance between the backbone atoms in the superimposed structures. To follow the dynamic evolution in the local components of the GQ structure, we calculated the RMSD individually for each of the components: GQ core, GQ loops and GQ LP loop. As shown in Figure 3, a), the GQ core exhibits an RMSD below 1 Å indicative of a highly stable and well-structured core. The dynamics of the G-A sheared base pair observed in the LP loop are demonstrated to be highly stable as well, since its RMSD value does not exceed 1 Åİn contrast, the single nucleotide loops have RMSDs fluctuating between 4 and 7 Å This behaviour is to be expected, given that these flexible free nucleotides are not initially engaged in a specific type of interaction. The root mean square fluctuations (RMSF) were also calculated to identify the most dynamic *c-kit* GQ bases. These were found to be the single nucleotides A1, A5, C9, and the dinucleotide C11-T12 (Figure 3, b). The flexibility of these bases significantly impacts the overall GQ dynamics.

**Figure 3:**
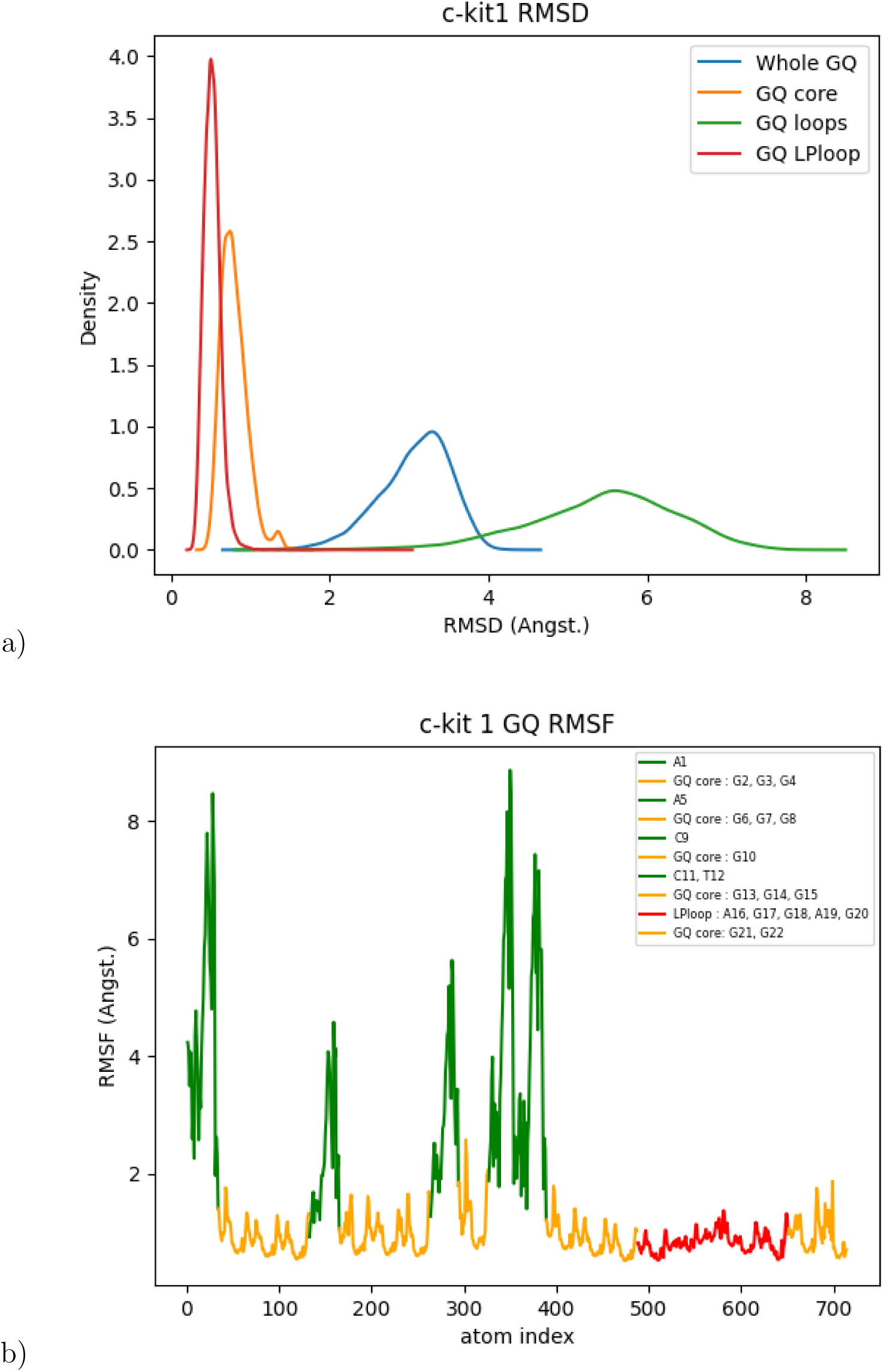
RMSD and RMSF of *c-kit1*. For the RMSF, the atom indices of nucleotides A1, A5, C9, C11 and T12 are respectively : 0 to 33, 133 to 164, 133 to 164, 133 to 164 and 133 to 164.

A better understanding of the dynamics of these nucleotides is enabled by the corresponding temporal evolutions of two torsion angles, four distances and of the *pi*–*pi* stacking indices as reported below.

#### Evolution of the loop conformations

We then investigated the evolution of the loop conformations. To do so, we monitored the *χ* glycosidic torsion angle evolutions, with the possible occurrence of *(anti / syn)* base flipping. The second monitored torsional angle is the angle *γ* governing the orientation of the sugar with respect to the phosphate backbone. Figure S1 and Figure S2 in the SI present the probability density of the *χ* and *γ* angles of A1, A5, C9, C11, and T12. C9 and C11 display large amplitudes of *χ* and *γ* values compared to A1, A5, and T12. A1 and T12 have limited *χ* angle variations (-50*^◦^* – +50*^◦^* and +25*^◦^* – +50*^◦^* respectively) but have a broad *γ* angle distribution. Such distributions enable the formation of a transient Hoogsteen basepair above the first GQ involving G2, G6, G10, and G13. It is stabilized by two H-bonds, between HN6(A1) and O4(T12), and between N7(A1) and HN3(T12). The involvement of N7 of A1 instead of N1 as in a standard W-C base-pair was unanticipated and appears to have no precedent in other GQ simulations. It locates N7 of A1 in the same groove as the three G bases, G2, G3, and G4 on the same strand which form the three tetrads. By contrast, the formation of a W-C base-pair was previously reported in the AMBER MD simulations. This Conventional Molecular Dynamics (CMD) simulation gave rise to the formation of a competing, longer-duration base-pair between A1 and C11. As an estimate, the time evolutions of the A1-C11 and A1-T12 base-pairs were monitored. We considered first the initial CMD trajectory (800 ns), then trajectories resulting from adaptive sampling (7.5 *µ*s). Figure S3 in the SI presents the CMD evolutions of the HN6–O4 distance for A1–T12 and of the HN6–N3 distance for A1–C11.

As shown in Figure 4, the initial shortening of the first distance to app. 2 Å is enabled by the flip movement of T12 with large amplitude *γ* fluctuations, and its stacking above the open face tetrad, particularly above G13 and G10. It initial occurs after 80 ns but only lasts 1 ns. It recurs again between 180 and 220 ns. At 260 ns, the formation of the concurrent A1-C11 base-pair takes place and this base-pair persists until the end of the CMD at 800ns. Such a non-classical interaction was not previously reported in polarizable force field nor in non-polarizable force field simulation studies.^12,30^ However, AMBER MD^12^ did record a movement of C11 over the helix terminal along with a short-lived interaction with either T12 or the phosphate backbone of G10. C9 and A5 remain highly dynamic and do not interact with any particular GQ nucleotide because of their locations in single nucleotide lateral loops. Within the stable LP loop two stable interactions can be detected, between A16 and G20, and between G17 and A19. We have, for both base-pairs, monitored the time evolution of one relevant intermolecular distance, between A16(N1) and G20(HN1), and between G17(HN2)–A19(N1). Such distances are subject to only limited variations around 2 Å for A16–G20 and 2.15 Å for G17–A19, attesting to the stability of the LP loop. Their onset occurs after 10ns. Somewhat larger amplitudes of variations, between 2 and 3 Å were reported in AMBER MD simulations.^12^ This result suggests that AMOEBA is able to detect and maintain Hoogsteen interactions between nucleic bases. Their onset starts after 10ns. Both AMOEBA and AMBER FF^12^ predict an additional GQ folding, which occurs through A1-C11 and A1-T12 H-bond interactions thus enhancing the overall stability of the DNA structure. These take place between one 6-NH2 hydrogen of A1 on the one hand, and both N3 of C11 and O4 of T12.A1 is found to interact alternatively with G2 and G13, while C11 and T12 interact with G6 and G10. Since the behavior of highly dynamic loops shows flip movements from the X-ray conformation and *π*-stacking interactions with the open face tetrad, we calculated the stacking index of the four bases in order to unravel the bases involved in the stacking. G2 is thus found to interact with A1 while both C11 and T12 interact with G10, Figure 7. Thus the stability of A1 above the open face tetrad could be favored by two types of interaction : a short-lived Hoogsten base-pair (*cis* with T12 and a long-lived *anti* WC base-pair with C11 together with a *π*-stacking interaction with G13.

**Figure 4:**
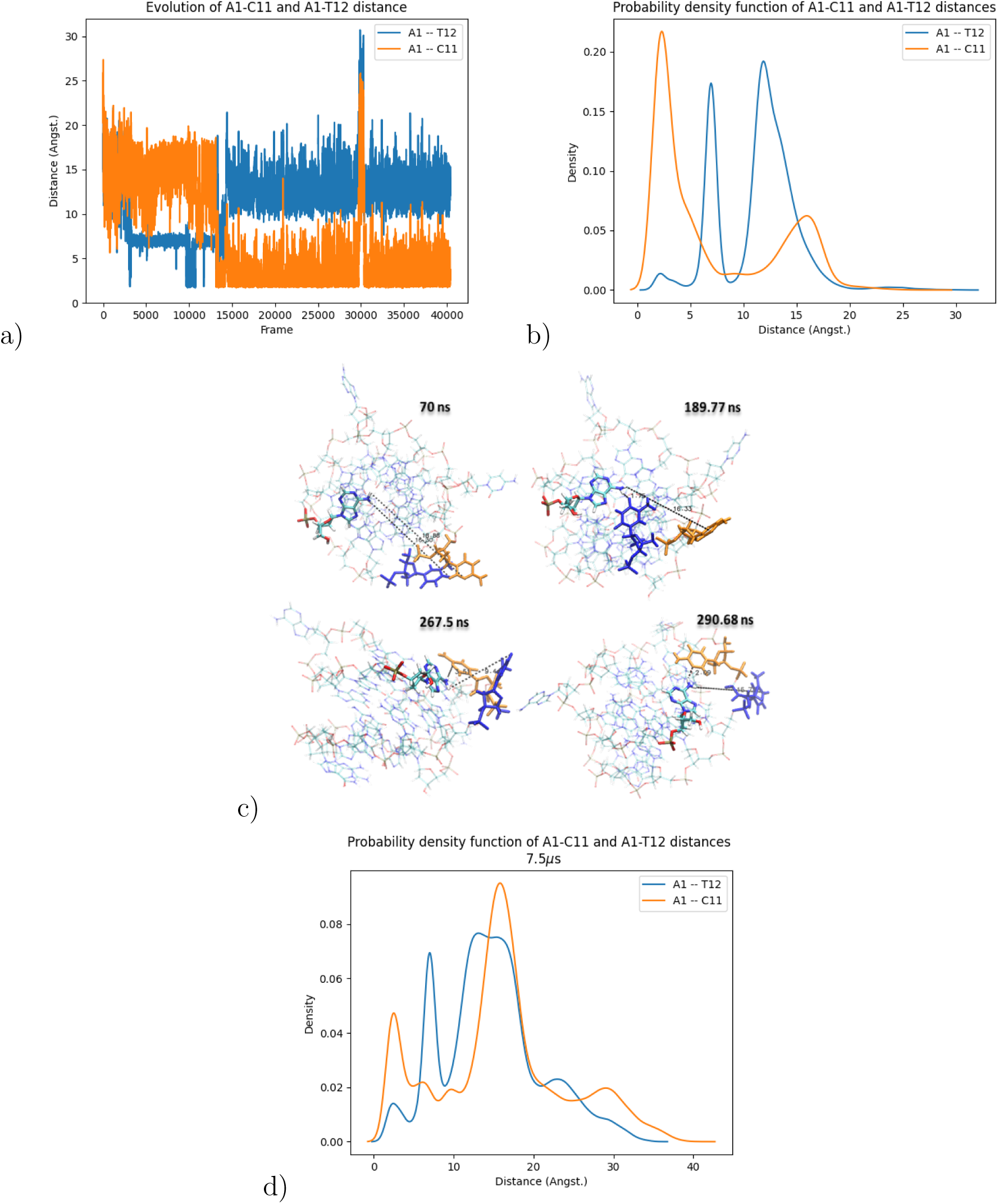
Evolution of the distance between A1(H) and T12(O) shown in blue and between A1(H) and C11(O) shown in orange. 5000 frames correspond to 100 ns simulation time. c) Representation of four structures at four different times of the CMD trajectory showing the evolution of distances A1–T12 and A1–C11.

### tICA clustering and cluster analyses

tICA is a dimensionality reduction technique^44–47^ designed to identify the slowest degrees of freedom (those with the highest autocorrelation) within trajectories. It facilitates the identification of microstates characteristic of molecular dynamics (MD) trajectories resulting from the unsupervised adaptive sampling strategy. These microstates are subsequently grouped into macrostates, each representing a distinct cluster. As described in our previous work,^35^ DBSCAN^48^ is specifically designed to identify clusters of arbitrary shape. We utilized two parameters: *ɛ*, the distance threshold at which two points are considered neighbors, and *MinPts*, the minimum number of neighboring points required to define a cluster. This clustering approach enables the characterization of conformational families, some of which may be targeted by rationally designed molecules.

Thus, we calculated the relative free energy of each cluster with respect to the most representative cluster (cluster 1). These energies were determined by evaluating the probability distribution across all sampled structures. As depicted in Figure 6 and Table 2 in the SI, cluster 1 exhibits the lowest free energy, indicating its highest probability of occurrence. Cluster 5 ranks second, with a free energy less than 1 kcal/mol higher, while the remaining clusters possess free energies 1.5-2.5 kcal/mol higher. The root mean square fluctuations (RMSFs) of the seven clusters were analyzed to quantify their dynamic behavior within the GQ. Figure 3 (SI) illustrates that nucleotide A1, single nucleotide loops A5 and C9, and the dinucleotide loop C11-T12 exhibit substantial fluctuations. With the exception of A5, these fluctuations demonstrate varying amplitudes across the clusters, ranging from 6-9 Å. In contrast, the LP loop and all twelve guanine bases of the GQ core display variations less than 2 Å. A1, A5, C9, C11, and T12 were therefore identified by the RMSFs of the tICA-derived clusters as the most flexible nucleotides of the *c-kit* GQ, contributing to the distinct conformational shapes of each cluster. To further characterize these clusters, we examined the evolution of the *χ* and *γ* torsional angles. Regarding the *χ* angles, Figure S1 in the SI demonstrates that for A1, A5, C9, and T12, all clusters exhibit the highest probability densities at similar values. However, for C11, the *χ* density in cluster 1 is centered around a Gaussian function peaking at approximately 10*^◦^*. In the other clusters, the *χ* density is centered around two major states, with peaks in the -120*^◦^* to -170*^◦^* and 120*^◦^* to 170*^◦^* regions. Concerning the *γ* angles, Figure S2 in the SI reveals that all clusters have their highest probability densities for A1 and A5 centered around -25*^◦^* and -110*^◦^*, respectively. Significant differences between clusters are observed for C9, C11, and T12. The highest probability density for the C9 *γ* angle is located between 75*^◦^* and 130*^◦^* for clusters 1, 3, 6, and 7, while clusters 2, 4, and 5 have their density peak at -40*^◦^*. C11 is highly structured in cluster 6, but in cluster 1, its *γ* angle exhibits two peaks, around -25*^◦^* and 80*^◦^*. In the remaining clusters, the *γ* probability density demonstrates shallow variations in the -100*^◦^* to -180*^◦^* and 100*^◦^* to 180*^◦^* regions. For T12, the *γ* peaks are around -50*^◦^* for clusters 1, 2, 4, 5, and 7, and around 60*^◦^* for clusters 3 and 6. These results indicate that C11 can undergo changes in its *χ* angle across the different clusters, enabling it to populate both *syn* and *anti* states. In contrast, A1, A5, C9, and T12 exhibit *χ* angles centered around a single peak in all clusters. Regarding *γ*, both A1 and A5 show values centered around a single peak in all clusters. C9 and T12 display two predominant values, centered around -40*^◦^* and 75*^◦^*. Finally, C11 exhibits one peak in cluster 6 and two peaks in all other clusters. These observations demonstrate the high stability of the GQ core and the LP loop in the *c-kit* GQ structure across all defined clusters. However, the single nucleotide loops (A5 and C9), the terminal nucleotide (A1), and the dinucleotide loop (C11 and T12) exhibit high flexibility, as they are not involved in stable intramolecular interactions.

**Figure 5:**
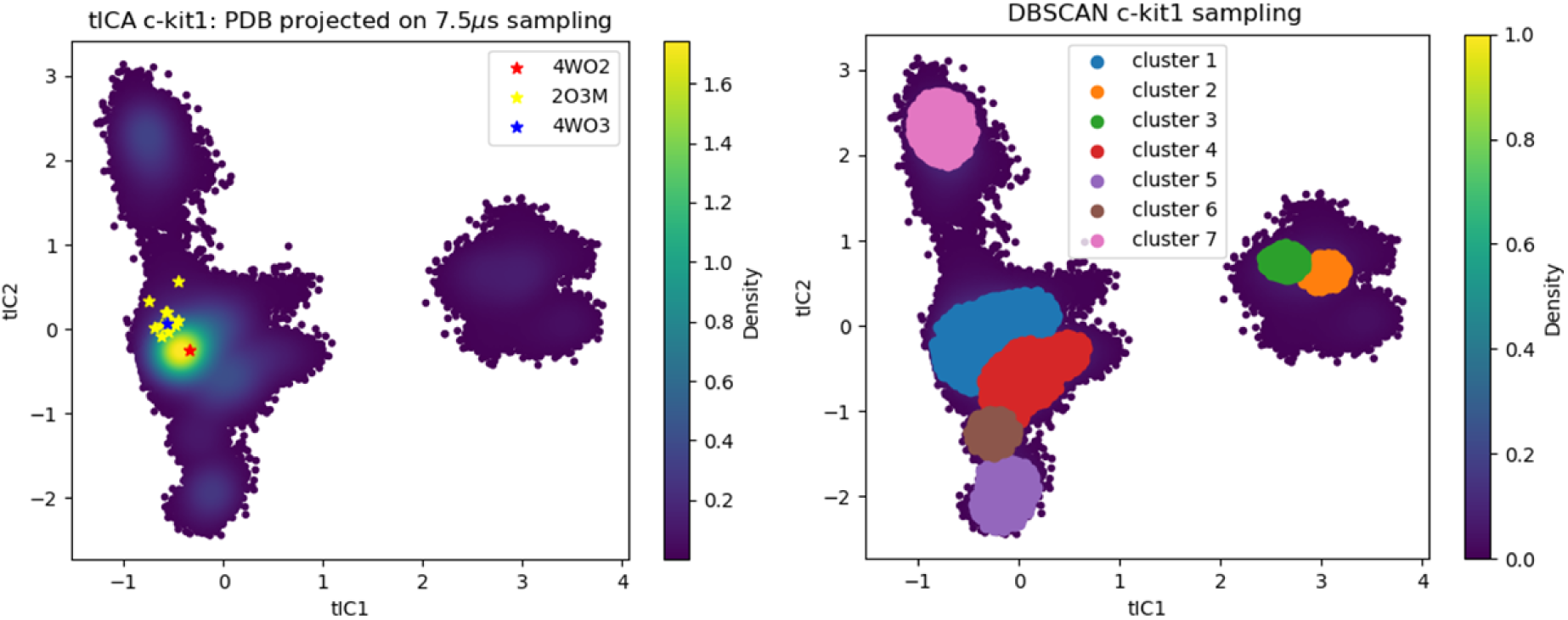
tICA DBSCAN sampling and clustering

**Figure 6:**
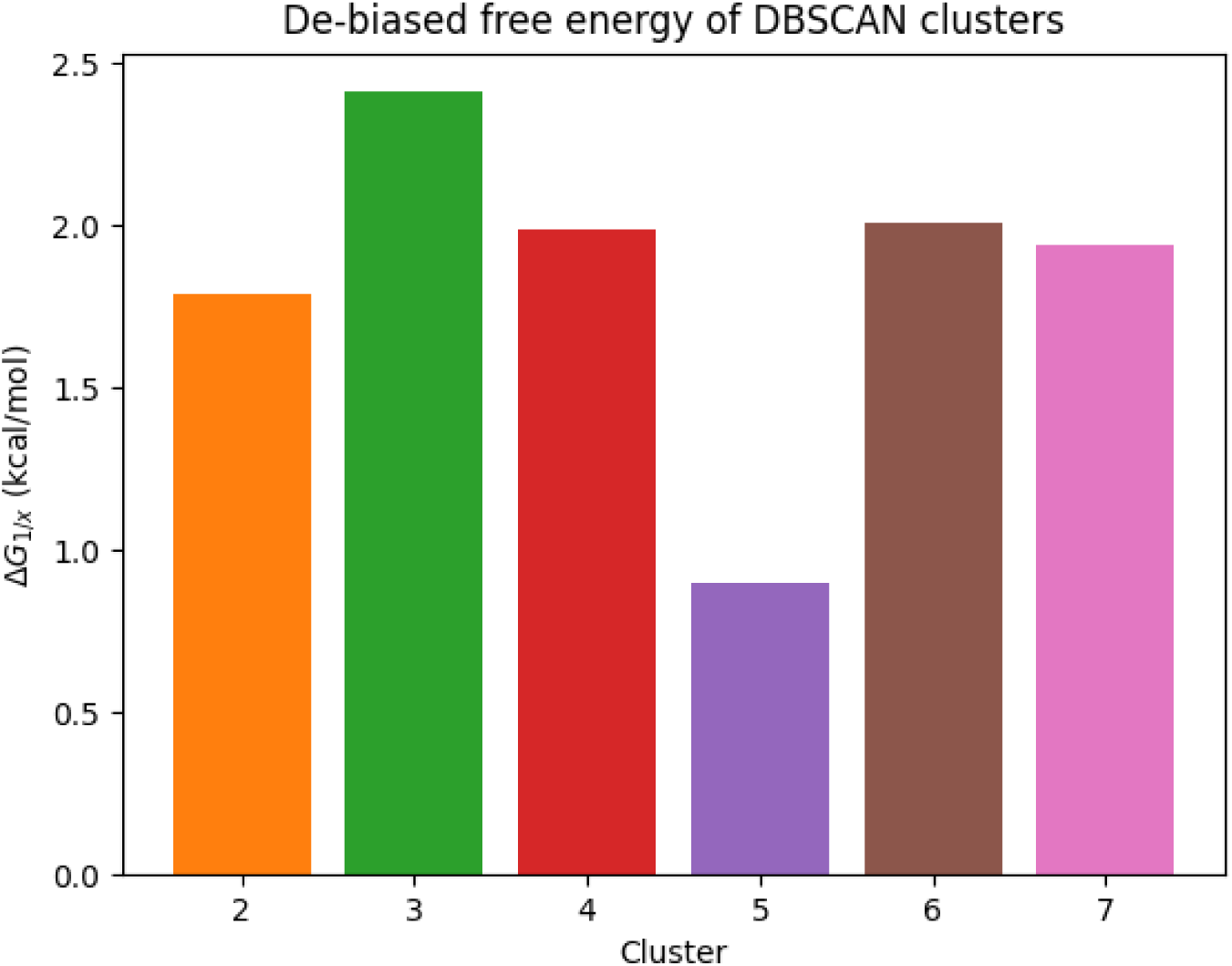
Relative free energies of the DBSCAN 7.5*µ*s tICA clusters, with respect to cluster 1.

However, both conventional and adaptative MD have identified transient H-bonding interactions between N1 of A1 and HN3 of C11. H-bond interactions were also detected between N1 of A1 and HN3 of T12, but are not maintained through the full time of MD simulations. The H-bonding distances involved were calculated accordingly for all the generated data (combining all the clusters). Figure 4 in the SI clearly shows that the highest probability density is associated to a defined H-bond interaction between A1 and C11, whereas the interaction between A1 and T12 is not as well characterized. Additionally, we have recently resorted to the *π*-*π* stacking index^49^ in an analysis of the M*^pro^* SARS–CoV–2 protein. It had enabled to quantify the extent of stacking between two residues, Phe140 and His163, a key interaction that stabilizes an oxyanion hole structure upon which the activity of each M*^pro^* protomer depends.^35^ Such an analysis is pertinent in the present study, as it can similarly evaluate the stability of the stacking of A1, C11, and T12 above the first guanine tetrad. In the result of the adaptive MD simulation, Figure 7, a), shows that A1 interacts by *π π* interaction with G13 in persistent mode. This is shown by a relatively high probability density for a stacking index of 0.2 comparing to an index of 0.15 for A1–G2 *π π* interaction. Figures 7, b) and c) show that C11 and T12 stack above G10 preferentially than above G6. Thus on the one hand, the stacking index for C11–G10 of 0.15 has a relatively higher probability density than the stacking index of 0.12 of C11–G6. On the other hand, the stacking index for T12–G6 is 0.10, smaller than for T12–G10 (0.18). The rearrangement of nucleotides A1, C11, and T12 to form a new plane of nucleotide interactions could further enhance the stability of *c-kit1* GQ. The formation of a hydrogen bonded complex between A1 and C11 esults in an improved *π* stacking index with respect to the guanines forming the first plane of guanine tetrads.

**Figure 7:**
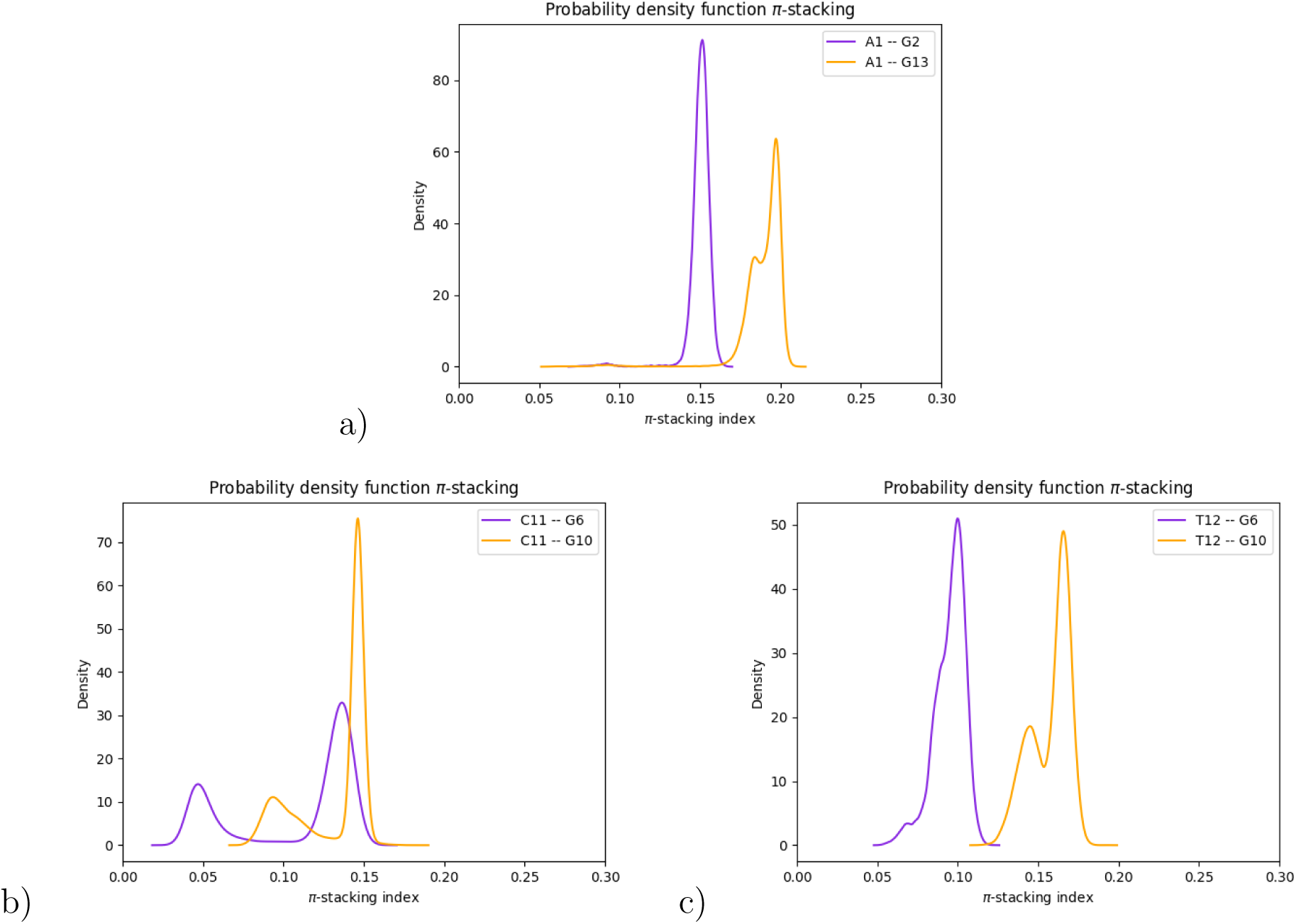
*π*-*π* stacking indexes for base pairs A1–G2, A1–G13, C11–G6, C11–G10, T12–G6 and T12–G10 calculated from the adaptive MD trajectories.

### GQ stabilization by K^+^ and polarizable H_2_O molecules

GQ interact with ions in an extremely dynamic fashion, causing experimental detection difficulties.^50^ Nevertheless, based on structural techniques,^51^ it was shown that tightly bound waters to G-quadruplex DNA are “NMR visible” in solution. In that context, it was put forth that K^+^ cations inside the channel exert an essential stabilizing role to the *c-kit* promoter and, indeed, high-resolution structures in the presence of K^+^ are available from 3D-NMR spectroscopy and X-ray crystallography. As discussed, we retained as a starting structure the crystal structure with the best crystallographic resolution (PDB: 4WO2^22^) for the 22-mer structure (1.82 Å) of the human native c-Kit proto-oncogene promoter. In order to study *c-kit* GQ behavior in the absence of these K^+^ cations, we performed 20 independent simulations of 20 ns each, using a similar setup as described in the Appendix, starting from an empty ionic channel where we displaced the structural K^+^ coordinates outside of the GQ structure to join the bulk ions.

We assessed the stability of the GQ structure by calculating its RMSD in each trajectory. Figure S5 in the SI shows that the *c-kit* GQ preserved its stability and initial conformation during all 20 trajectories, with some minor conformational changes of the terminal nucleotide (A1) and dinucleotide loop (C11 and T12). GQ core stability can be explained by the fact that the ionic channel was never seen empty by the end of any of these MD simulations. We counted the number of H_2_O molecules and K^+^ cations inside the ionic channel for every frame of these 20 MD simulations to understand the process of the ionic channel filling. To do so, we defined a virtual sphere centered on the mass center of the 12 guanines contributing to the formation of the *c-kit* GQ core and with a radius of 4 Å (which is the best radius to cover all and only the ionic channel). As it is shown in Figure 8, and in more detail in Figure S6 in the SI, several scenarios of ionic channel filling could be put forth. The highest density detected is associated with the presence of 1 H_2_O molecule and 1 K^+^ cation inside the ionic channel. The second highest density corresponds to an empty channel where no H_2_O molecules, nor cations, are detected inside the GQ core. No channel was found empty at the end of any of the 20 MD simulations, and the density here corresponds to frames from the beginning of the trajectories in which the ionic channel is empty. The third highest density observed corresponds to 2 H_2_O molecules and 1 K^+^ cation that can be present inside the ionic channel. With a slightly lower density, we can observe either the presence of 2 H_2_O molecules, or that only 1 H_2_O molecule only.

**Figure 8:**
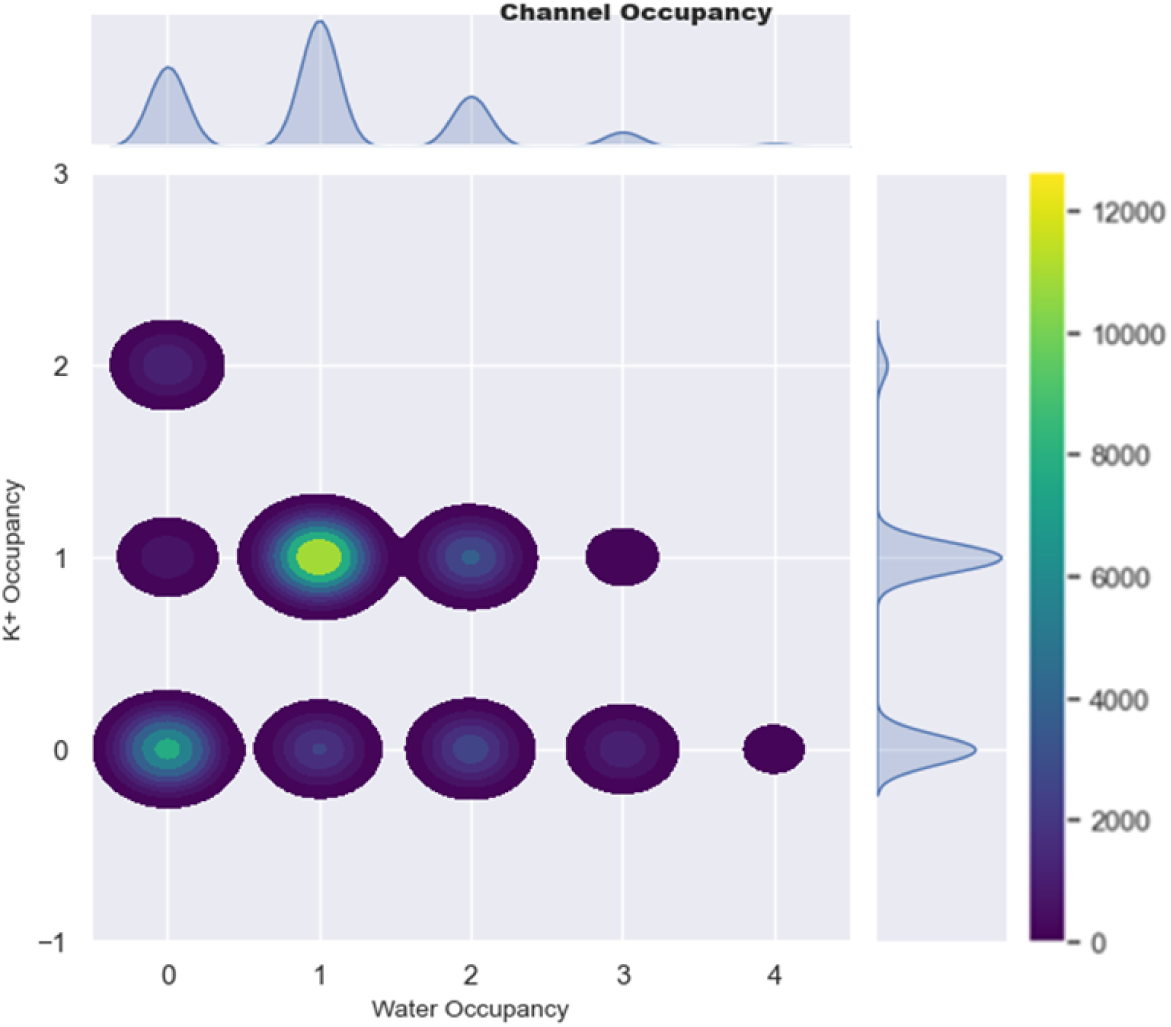
2D probability density function representation of *c-kit1* GQ channel occupancy throughout the 20 independant trajectories of 20ns each starting from an empty channel. The color bar represents the frame density, from blue to yellow we represent respectively the lowest and the highest density region.

Once the channel is filled, all loops evolve in a similar fashion as in the MD simulations where the two K^+^ cations were present in the channel since the start of the simulation with no significant relative structural changes. Over the 800 ns of accumulated CMD trajectory, Figure S7 (Supporting Information) demonstrates that both K^+^ ions, present from the start of the simulation, remained within the *c-kit1* GQ channel, with no water molecules observed inside.

Previous molecular dynamics (MD) simulations ^52^ have indicated that potassium ions (K^+^) at the terminus of a GQ structure (PDB ID 1JRN) referring to an orthorhombic form of an orthorhombic form of Oxytricha telomeric DNA, can readily dissociate within the initial stages of simulation. To further validate this hypothesis, we performed on this structure a 50ns MD simulation, observing a consistent retention of K^+^ within the GQ channel as seen in the aforementioned study, where K27, initially positioned within the channel, rapidly dissociates. Subsequently, K25 and K28 ions migrate to occupy the vacated site. The distance from the original position of each of the K^+^ was calculated and presented in Figure S12, a). Structurally, and as shown in Figure S12, b) in the SI, K27 and K26 are positioned between guanine tetrads, surrounded by a thymine loop, which provides the terminus with high solvent accessibility. This suggests that the observed stability of K^+^ within the GQ channel may be attributed to the specific DNA conformation and the presence of guanine tetrads sandwiching the cations where the cation stabilization within the channel is facilitated by an equilibrated arrangement of oxygen atoms from the guanines pointing inwards.

We calculated the average dipole moments of the H_2_O molecules that have entered the channel. Their mean value (2.72 Debye) is slightly below the one of the AMOEBA bulk water (2.78 Debye). This, albeit modest, decrease could possibly be due to a local decrease of the polarizing guanine-induced field inside the channel, due to the mutually-opposing fields exerted in each tetrad by the two pairs of two symmetry-related guanines. In this connection, it was recently observed^36^ that a dense and compact environment generates a global many-body depolarizing effect influencing the H_2_O molecule induced dipoles.

It was found in G4-wire^53^ simulations that several water molecules progressively fill the G-channel during the dynamics.CHARMM simulations resulted into one K^+^ from the solvent phase entering the channel after 20 ns.^30^ This entrance was from below tetrad 3 rather than, as in AMOEBA MD, from above tetrad 1. It reached tetrad 3 after transiting through the Long Propeller Loop, which, as mentioned in,^30^ could cause transient conformational distortions in the LP loop and tetrad 3. Drude simulations, ^30^ starting from an empty channel, resulted into one K^+^ transiently interacting with T12 and then entering the GQ core through the first tetrad, as found here. The 20 AMOEBA MD simulations run in parallel, started from an empty channel, showed the influx of three or four water molecules to occur shortly after the beginning of the MD simulations. These could buffer the electrostatic repulsions between the C=O groups, providing the requisite time for K^+^ to enter the channel. The subsequent entry of one K^+^ is synchronous with the exit of one or two water molecules.

A previous non-polarizable MD simulation focused on a pre-folded GQ structure, the thrombin-binding aptamer 15-TBA.^54^ In a water solution without cations inside the GQ channel, its structure was found stable for a sufficiently long time to enable eventual capture of a cation from the bulk. Furthermore, ion exchange events were observed without any GQ core destabilization.^55^ On the other hand, the instability of cation-depleted GQ channels was reported in previous publications.^53,56^ This could possibly be ascribed to the absence of explicit polarization effects and, as a consequence, the lack of local cooperativity, which would have further buffered the mutual repulsion between the C=O groups. We did not attempt simulations devoid of K^+^ cations to evaluate the lifetime of a water-filled, cation-depleted GQ channel, as this is not the objective of this work. It is noted that the positions of the water molecules within the channel do not precisely align with that of the K^+^, but rather exhibit a varying degree of coordination contingent on the number of water molecules present and the presence or absence of a K^+^.

It was previously reported^56,57^ that with some non-polarizable force fields, certain GQ geometries could be stabilized by ‘bifurcated’ hydrogen bonding interactions, in which N1 donates its proton to both O6 and N7 of one neighboring guanine, while N2 donates a proton to N7 of that neighboring guanine. We verified the possibility of observing a bifurcated geometry between all guanines in the GQ tetrads for AMOEBA MD simulations. Figure S8 in the supplementary information (SI) illustrates the probability density distribution of the distances between the atom N7 and the atom H1 of every two adjacent guanines within a single tetrad. It can be observed that the distance between these atoms is never sufficiently close to indicate the onset of bifurcated hydrogen bonding.

### Cryptic pockets

We used the DoGSite Scorer software^58^ to identify potential drug binding cavities in each of the seven identified clusters. For all seven clusters, two pockets were defined: P1, formed by bases A1, G2, G3, G13, G14, G15, A16, and P2, formed by bases G7, G8, C9, G10, C11, G21, as shown in Figure 9. To further investigate how key pocket inter-base distances persist throughout adaptive MD, and how their values compare across the seven clusters, we considered three pairs within P1: G3–G15, G2–G14, and A1–G13, and two pairs within P2: G21–G8 and G10–G7. In the G–G pairs, the first G belongs to one tetrad and the second to the subsequent tetrad. It is stacked below the guanine that donates two hydrogen bonds to the first G. The intermolecular distances monitored are between N7 of the first G and HN1 of the second. In the A–G pair, the corresponding distance is between N7 of A1 and HN1 of G13, which is part of the first tetrad. This choice of G–G distances should enable evaluation of whether partial unstacking of the quadruplexes or distortion of the GQ hydrogen bonding occurred during MD. The A1–G13 distance exhibits large fluctuations in all clusters, particularly in clusters 5 and 7. This is clearly attributed to the large-amplitude motion of A1, whose RMSF exceeds 9 Å in these two clusters. However, Figure S11 in the SI shows that in all seven clusters, for each G–G pair, these distances remain closely distributed around sharp peaks within a 4-4.5 Å interval. K^+^ cations remain in place and coordinate the guanines within the GQ core, as shown in Figure 9 in the SI. This attests to the stability of the GQ core in all clusters. It should favor the binding of an incoming ligand to the groove formed by the bases belonging to P1 or P2.

**Figure 9:**
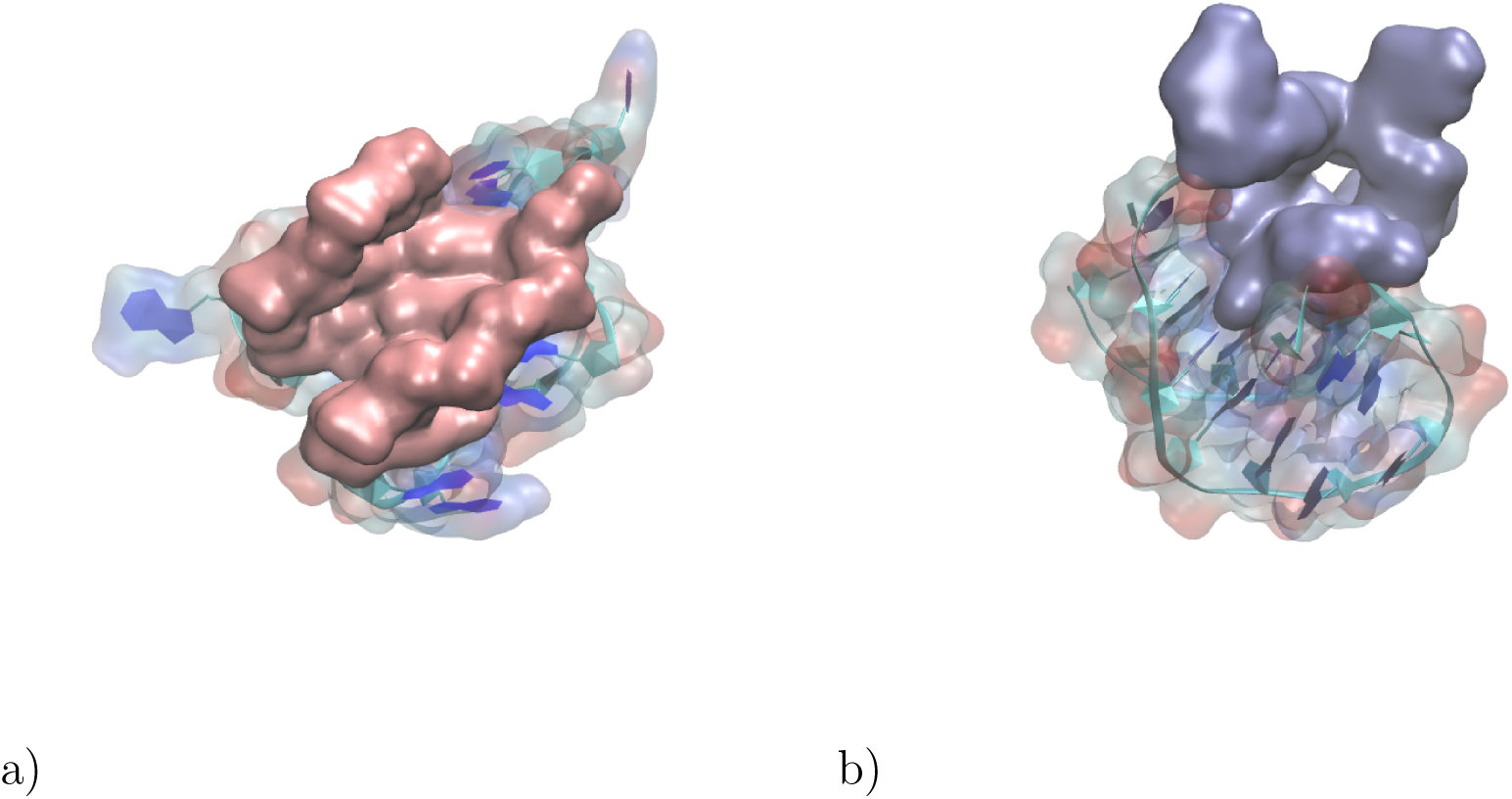
Representation of the two cryptic pockets inside *c-kit1* GQ; a) P1 (pink color) formed by the bases A1, G2, G3, G13, G14, G15, A16, and b) P2 (ice blue) formed by bases G7, G8, C9, G10, C11, G21.

### Simulation of the LTR-III region of HIV

The HIV-1 Long Terminal Repeat (LTR) is a crucial regulatory region that controls viral transcription and replication. Given that this region is also prone to GQ formation, we performed a preliminary 400 ns CMD simulation using the AMOEBA force field, with a high-resolution NMR structure as a starting point (PDB ID: 6H1K). DoGSite Scorer identified two cryptic pockets: P1, formed by bases G2, A4, G5, G6, G8, T9, G11, C12, C13, T14, G15, and G19, and P2, formed by bases G1, G2, G3, G15, G19, G20, and G21. P1 is primarily located within the hairpin domain, while P2 is located on the upper face of the GQ domain (Figures 10 a and b). Similar to the *c-kit1* GQ, root mean square deviation (RMSD) and root mean square fluctuation (RMSF) calculations (Figure 10, a and b, respectively, in the SI) demonstrate a highly structured GQ core. In contrast, the hairpin domain and both the three-nucleotide and single-nucleotide loops are unstructured.

**Figure 10:**
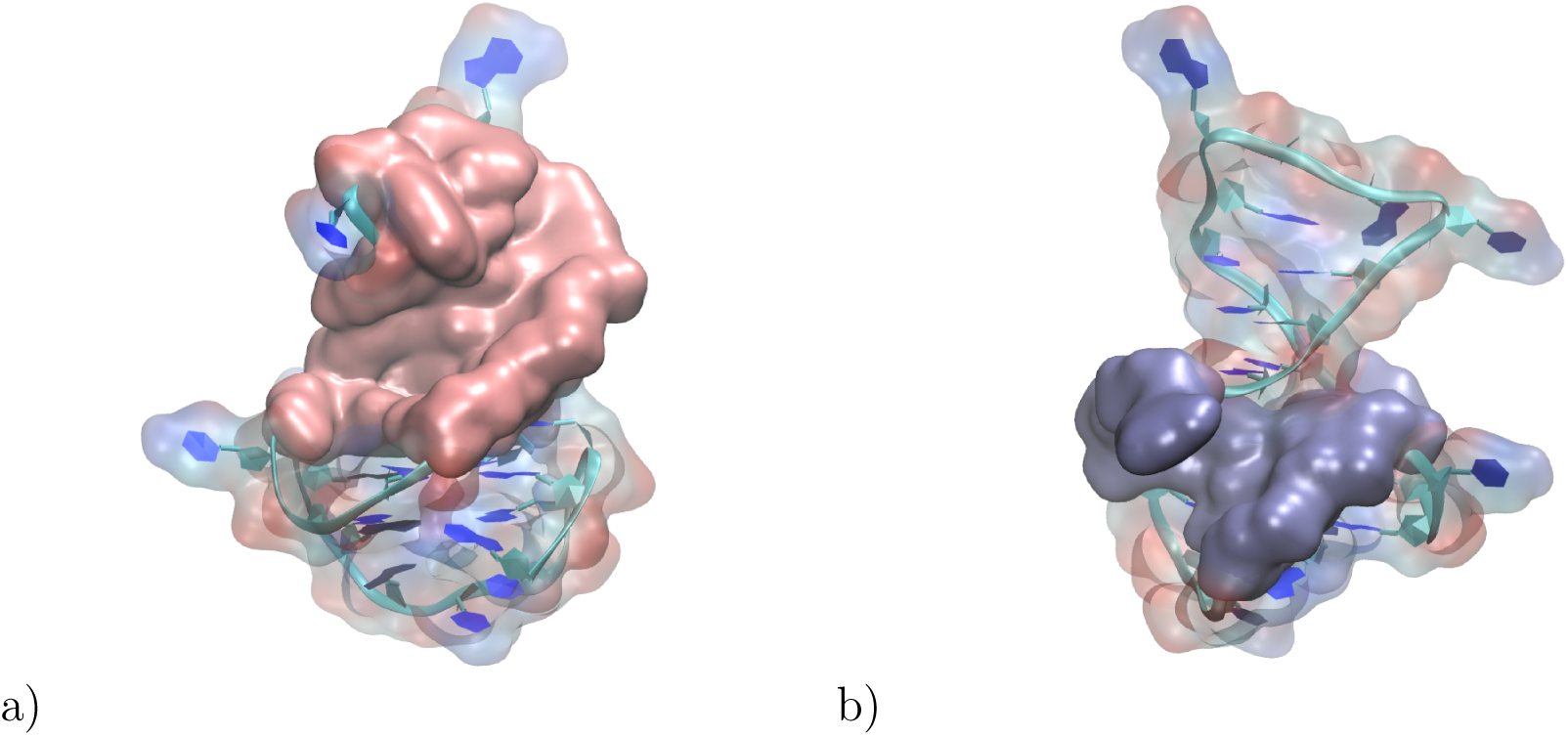
Representation of the two cryptic pockets inside LTR-III HIV GQ structure c) P1 (pink color) is formed by the hairpin residues and d) P2 (iceblue) is formed by the GQ core bases.

Previous MD simulations using Amber18^24^ reported similar RMSD and RMSF evolutions to the present AMOEBA simulations and also demonstrated a close agreement between simulation-derived and experimental data.

As reported above for *c-kit1*, we have monitored representative inter-base distances within P1 and P2. In P1, we have considered the three W-C base pairs A4–T14, G5–C13 and G6– C12. In P2, the two base pairs are G2–G19 and G1–G20. For the A–T base pair, the distances were measured between H3N(T) and N1(A). For the two G–C base-pairs, they are between H1N and N3. In P2, for both G2–G19 and G1–G20, they are between the O6 atoms of the two bases. This choice, rather than the one done for *c-kit1*, was motivated by the different glycosidic angles of the consecutive bases: anti for G2 and G20, and *syn* for G1 and G19. This results in the N7 atoms of two consecutive bases being located on opposite strands. The O6-O6 distances were considered less sensitive to the alternation of the glycosidic angles. By contrast, all GQ core guanines in*c-kit1* are anti.

The distance probability density functions (Figure 9c in the SI) indicate greater stability for P2 than P1. Specifically, the two O6-O6 distances in P2 are clearly centered around two peaks within a 4.5-4.8 Å interval. In P1, the A4–T14 distance exhibits two peaks. The first corresponds to a standard W-C hydrogen bond distance, while the second, at 4.2 Å, corresponds to a dissociated pair. Similarly, for G6–C12, there are two peaks, the first at 2.0 Å corresponding to a W-C base pair, and the second at 4.0 Å corresponding to a dissociated base pair. For G5–C13, a broad distribution is observed within the 2.0-4.5 Å range. Since the cryptic pocket P2 demonstrates greater conformational stability throughout the MD simulation, as evidenced by the distance probability density functions, drug design efforts should focus on targeting this substructure rather than pocket P1.

### Regarding the druggability of the guanine quadruplexes: from *c-kit1* to the LTR-III of HIV-1

As mentioned above, *ckit1* is a receptor tyrosine kinase involved in cell proliferation, survival and differentiation. Mutation or overexpression in *ckit1* are associated with cancers, notably gastrointestinal stromal tumors (GISTs). GQ have been indentified in the promoter region of *ckit1* gene and thus can influence gene expression making them potential targets for anti-cancer drug design. DoGSite Scorer identified two ‘cryptic’ pockets, P1 and P2, which could serve as ‘druggable’ sites for potential pharmacological ligands. Figure 9 a) gives a representation of the accessible surface of P1, clearly revealing the existence of a suitable for ligand binding.

Since GQs were also found within the HIV LTR-III region, a novel approach to anti-HIV therapy could be proposed. We have performed a 400 ns CMD on the GQ found on the LTR-III of HIV-1. Here also, DoGSite Scorer identified two cryptic pockets, P1 and P2, of which only P2, which exhibited more limited RMSD variations, could potentially serve as a druggable ligand binding site. Figure 10 provides a representation of the accessible pockets.

Based on these results, the design of novel ligands targeting the grooves of cryptic pockets P1 and P2 of *c-kit1*, and of P2 of the LTR-III of HIV-1 appears feasible. This approach could be extended to other oncogenic and retroviral sequences prone to G-quadruplex formation. The rationale for targeting GQs relies on their unique structural features, which make them attractive targets for drug design. Molecules that specifically bind to and stabilize GQs could potentially disrupt biological processes without affecting other cellular functions or causing significant side effects.

## Conclusions

We have performed long AMOEBA polarizable MD simulations on two guanine quadruplexes (GQs). The first is formed within an GC-rich oncogenic sequence, associated with the *c-kit1* oncogene, and the second is found within a Long Terminal Repeat of the HIV-1 retrovirus. We first focused on a 22-mer oligonucleotide belonging to the *c-kit1* oncogene, whose overexpression can lead to cancer. This oligonucleotide can fold into GQ, a conformational change occurring during DNA transcription into CD117 mRNA. Understanding the dynamics of this 22-mer oligomer in its quadruplex is essential and was analyzed in detail. Using the AMOEBA polarizable force field coupled to GPU-accelerated unsupervised adaptive sampling, we achieved an aggregated*c-kit1* simulation time of 7.5 *µ*s. Most simulation features were found to be fully consistent with those previously obtained using the non-polarizable AMBER^12^ or polarizable Drude ^30^ force fields. However, while the long-duration stability of the GQ core, composed of three stacked guanine tetrads, and that of the Long-Propeller Loop observed in previous studies were confirmed, this contrasts with the high dynamics of the single-nucleotide loops A5, C9 and of the dinucleotide loop C11-T12, whose mobilities impact the overall dynamics of the *c-kit1* 22-mer. Such mobilities were evaluated in the individual clusters in terms of: a) H-bonding distances in base-pairs A1-T12 and A1-C11, A16-G20, and G17-G20; b) the two torsion angles, i.e. the glycosidic *χ* and the *γ* torsion angles of bases A1, A5, C9, C11 and T12, and c) the *π*-stacking index between A1 and both bases, C11 and T12 of the dinucleotide loops, and a G base of the nearest guanine tetrad. We have also monitored the evolution in the course of adaptive MD for base-pairs A1-G2, A1-G13, C11-G6, C11-G10, T12-G6, and T12-G10. The tICA analyses revealed seven distinct clusters transiently stabilized during the MD simulations. Available X-ray structures were identified in the largest detected cluster. The fact that the identified clusters remain closely related in terms of energy is due to the fact that they are all almost identical when it comes to the strucutre of the GQ core and the LP loop. The difference between these clusters arises from the varying conformations of the single and dinucleotide loops. However, all of these clusters share the same stable cryptic pockets which can therefore be considered potentially druggable sites for drug discovery. GQ’s channel simulations using the AMOEBA polarizable force field demonstrated the importance of polarizable water molecules to transiently stabilize the channel against unfolding until K^+^ influx occurs. Our modeling strategy was extended to another model and thus was tested on the LTR-III region of HIV-1. The stability of the G-quadruplexes was also assessed. Well-defined and stable cryptic pockets were also detected. One of them can be proposed as a potential druggable site.

## Technical Appendix

### Materials and methods System construction

The starting structure for *c-kit1* GQ was taken from the crystallography structure in PDB entry *4WO2*.^23^ The structure was solvated in a water cubic box with dimensions of 60 Å. To mimic physiological conditions, 150 mM of [KCl] (K^+^, Cl^-^) was added to the system. The system was then minimized with the AMOEBA polarizable force field^31,59,60^ using L-BFGS algorithm until a Root Mean Square (RMS) gradient of 1 kcal/mol was reached.

### Conventional Molecular Dynamics

The MD simulations were performed using the recently developed GPU module^42^ of the Tinker-HP software,^43^ which is part of the Tinker platform.^61^ This module efficiently leverages mixed precision,^42^ offering a substantial acceleration of simulations. The assembly was equilibrated for 1ns at each of the following temperatures successively: 50, 100, 150, 200, 250, and 300K. The Smooth Particle Mesh Ewald^62^ was used with the Ewald and van der Waals cutoffs set to 7 Å and 9 Å, respectively. The MD simulations used the BAOAB-RESPA integrator with a 2 fs outer timestep, a Bussi thermostat, hydrogen-mass repartitioning (HMR), and random initial velocities. Smooth Particle Mesh Ewald (SPME) method was used for handling periodic boundary conditions with a cubic boox of dimensions 60 Å x 60 Å x 60 Å. The production phase was conducted at 300 K for 800 ns for*c-kit1* and 400 ns for HIV-1 LTR-III.

### Unsupervised Adaptive Sampling strategy for conformational exploration

Adaptive sampling has proven to be a powerful exploration tool for studying protein folding and dynamics, as well as for exploring a diversity of rare molecular events.^63–66^ Recently, we developed a new fully unsupervised Adaptive Sampling strategy that enables the simultaneous use of hundreds of GPU cards. ^35^ The method divides the simulation procedure into iterations, each composed of a swarm of independent MD simulations run in parallel.

At the beginning of each iteration, some initial structures are selected from the structures sampled in past iterations. The selection procedure is mathematically designed to fully explore a low-dimensional space of slow variables generated by dimensionality reduction algorithms such as PCA and tICA.

From these selected structures, independent MD simulations are run in parallel, potentially combined with other enhanced sampling methods,^67^ to generate new structures that can serve as starting points for future iterations.

In this particular case, we first ran an initial 800 ns of cMD where we selected 50 starting point structures using the Adaptive Sampling algorithm in the four-dimensional PCA space. The choice of 50 starting points is heuristic, as we have observed it to be sufficient for efficiently sampling the conformational space of a protein of similar size: the SARS–CoV–2 protein. The number of dimensions for the PCA space is determined based on the fact that the other PCA dimensions have negligible variance contribution below 4 %. Each cMD simulation sampled 15 ns, and these simulations were performed in the NVT ensemble at 300 K. Thus, with a total of 10 iterations, we generated more than 7.5 *µ*s of simulation.”

### Time-structure Independent Component Analysis

Time-structure Independent Component Analysis (tICA) was introduced into MD as a method to extract slow collective variables.^44,45^ tICA solves a variational approach by which it finds an optimal approximation to the eigenvalues and eigenfunctions of the Markov operator underlying the molecular dynamics data. ^44,46,47^ As a result, tICA is ideally suited for clustering and to prepare data for Markov models.

In fact, tICA followed by clustering techniques makes it possible to classify the structural and physico-chemical properties of the structures within an MD trajectory. Additionally, this approach reveals valuable insights into the system’s behavior over time.

While tICA shares similarities with PCA, it offers notable advantages for capturing the dynamics of proteins. Unlike PCA, which relies on static and linear properties, tICA aims to create independent components as meaningful as possible to better describe the dynamics of a system.^68^ This makes tICA a more suitable choice for our analysis.

Throughout the simulations, with tICA and clustering techniques we identified distinct clusters for each of the variants analyzed. We organized the final MD trajectories into clusters based on the eigenvalues and eigenfunctions of the covariance matrix.

By employing this classification approach, we can observe similarities between the macrostates of each variant, enabling us to discern common structural or dynamic changes shared among the variants.

A classification makes it possible to observe the similarities between the macro-states of each of the variants to discern a structural or dynamic change to them.

## Acknowledgement

DEA thanks funding from the Lebanese National Council for Scientific Research, CNRS-L. FC thanks funding from the French state funds managed by the CalSimLab LABEX and the ANR within the Investissements d’Avenir program (reference ANR11-IDEX-0004-02) and support from the Direction Génerale de l’Armement (DGA) Mâıtrise NRBC of the French Ministry of Defense. This work was also made possible thanks to funding from the European Research Council (ERC) under the European Union’s Horizon 2020 research and innovation program (grant agreement No 810367), project EMC2, JPP). Computations have been performed at GENCI (IDRIS, Orsay, France and TGCC, Bruyères le Chatel) on grant no A0130712052 (JPP).

## Author Contributions

DEA contributed to conceptualization, methodology, investigation, formal analysis, validation, visualization and writing. LL contributed to investigation, software support and writing. TJI contributed to data curation and methodology. FC contributed to conceptualization and writing. ZH, NG, RM and JJP contributed to conceptualization, supervision, validation and writing. RM and JPP contributed to funding acquisition.

The authors declare no competing financial interests.

## Data and Software Availability

Calculations were performed using the Tinker-HP software (https://www.tinker-hp.org/) which is freely available for academics on GitHub (https://github.com/TinkerTools/tinker-hp). All simulation parameters were described in the Technical Appendix section, and AMOEBA force field parameters were taken from the published literature, see references.

## Supporting Information Available

Supplementary information is available. It includes: the free energy values of each of the identified tICA clusters; the nucleotides probability density functions; the RMSF of ckit1 GQ components in each of identified tICA clusters.; the*c-kit1* GQ RMSD for each of the 20 independent MD simulations starting from an empty ionic channel.

