## Supplementary Information for "AMOEBA Polarizable Molecular Dynamics Simulations of Guanine Quadruplexes: from the c-Kit Proto-oncogene to HIV-1"

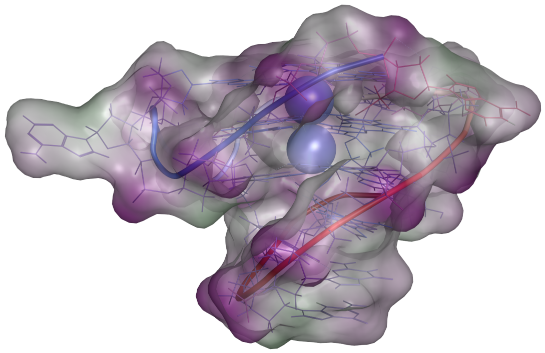

Table S1: Molecular dynamics simulation summary

| | |
|---|---|
| *c-kit1* GQ CMD | 800 ns |
| *c-kit1* GQ Adaptive Sampling | 7.5 $\mu$s |
| *c-kit1* GQ CMD starting from emtpy channel | $20 \times 20$ ns |
| LTR-III HIV GQ CMD | 400 ns |
| PDB ID 1JRN GQ CMD | 50 ns |

Table S2: Free energy values of each of the identified tICA clusters.

| (Kcal/mol) | Free Energy | $\Delta G$ with respect to cluster 1 |
|---|---|---|
| Cluster 1 | 0.57 | 0 |
| Cluster 2 | 2.36 | 1.79 |
| Cluster 3 | 2.98 | 2.41 |
| Cluster 4 | 2.55 | 1.98 |
| Cluster 5 | 1.46 | 0.89 |
| Cluster 6 | 2.58 | 2.01 |
| Cluster 7 | 2.51 | 1.93 |

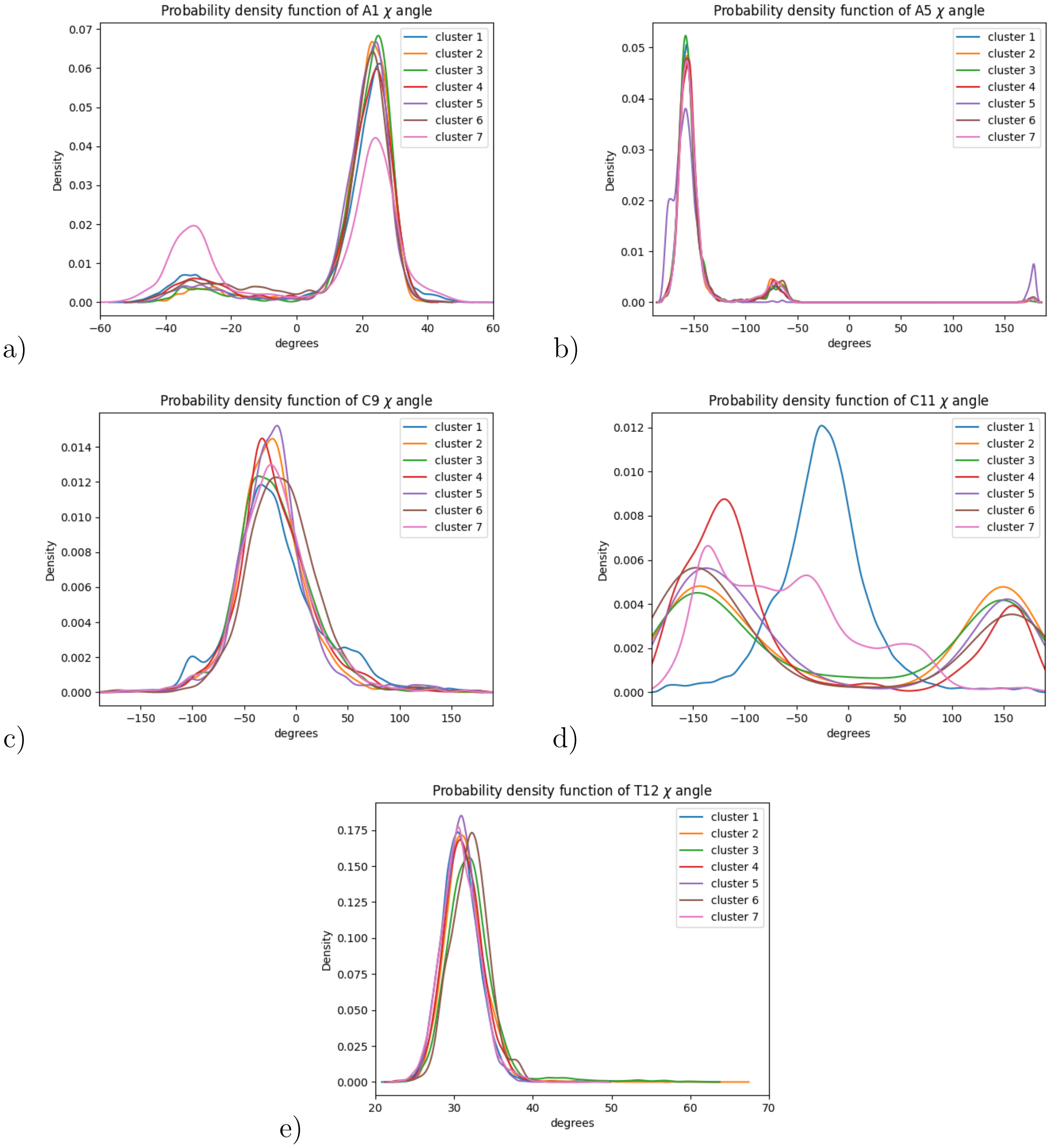


Figure S1: Probability density functions of A1, A5, C9, C11 and T12 nucleotide $\chi$ angle in the seven tICA clusters, respectively in a), b), c), d) and e).

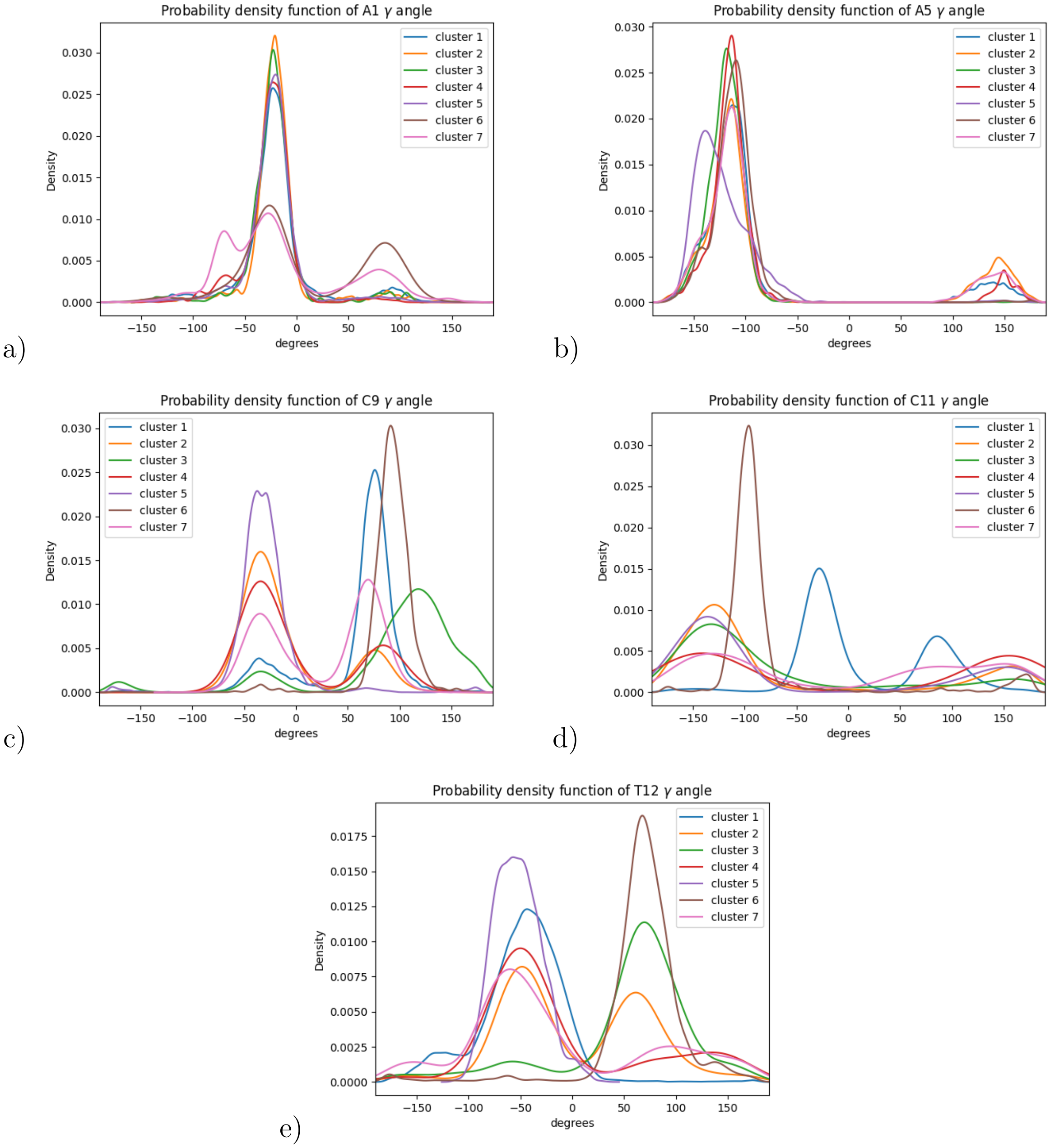


Figure S2: Probability density functions of A1, A5, C9, C11 and T12 nucleotide $\gamma$ angle in the seven tICA clusters, respectively in a), b), c), d) and e).

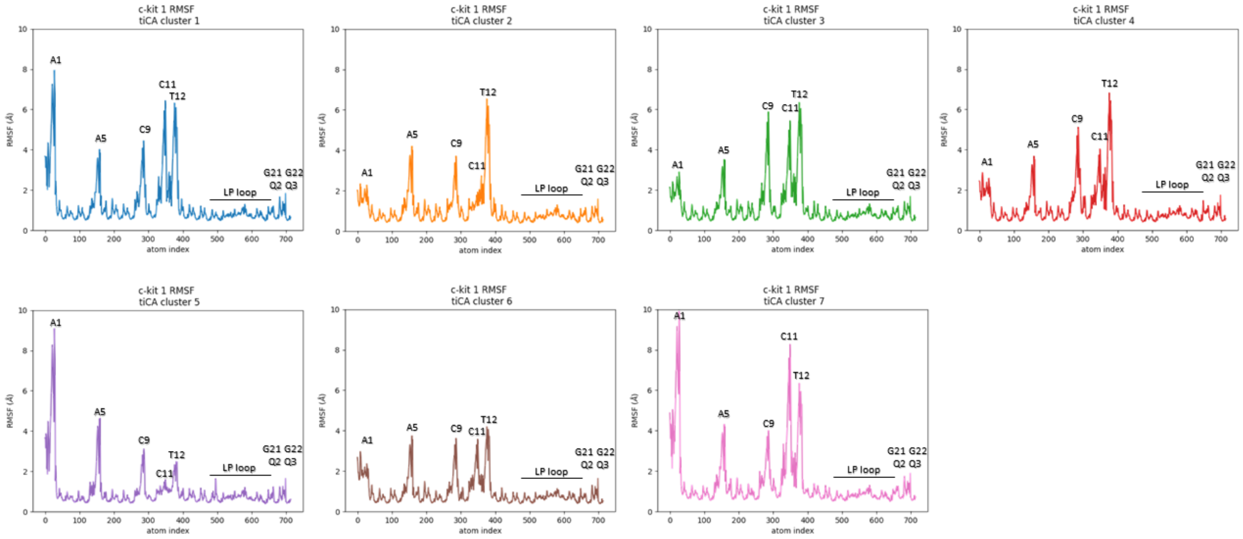


Figure S3: RMSF of *ckit1* GQ components in each of identified tICA clusters.

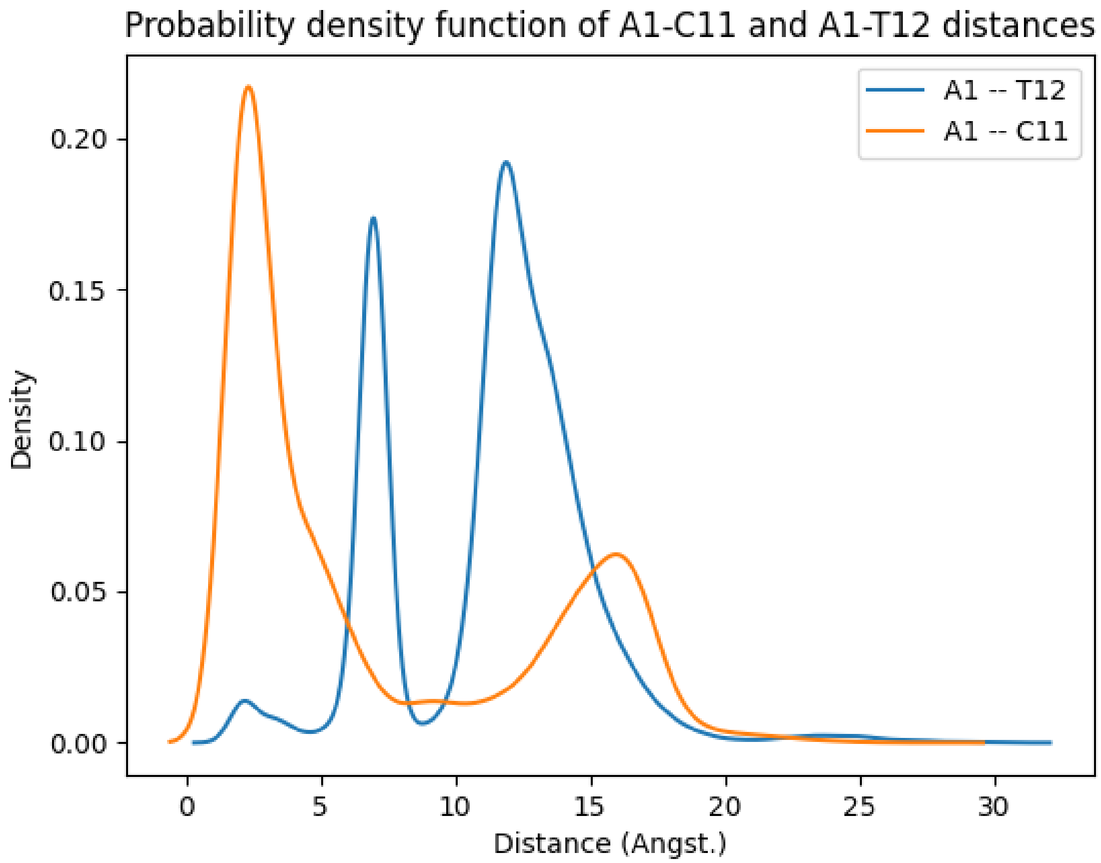


Figure S4: Probability distribution of the distances between A1 and C11 and between A1 and T12 for all the generated data (combining all the clusters).

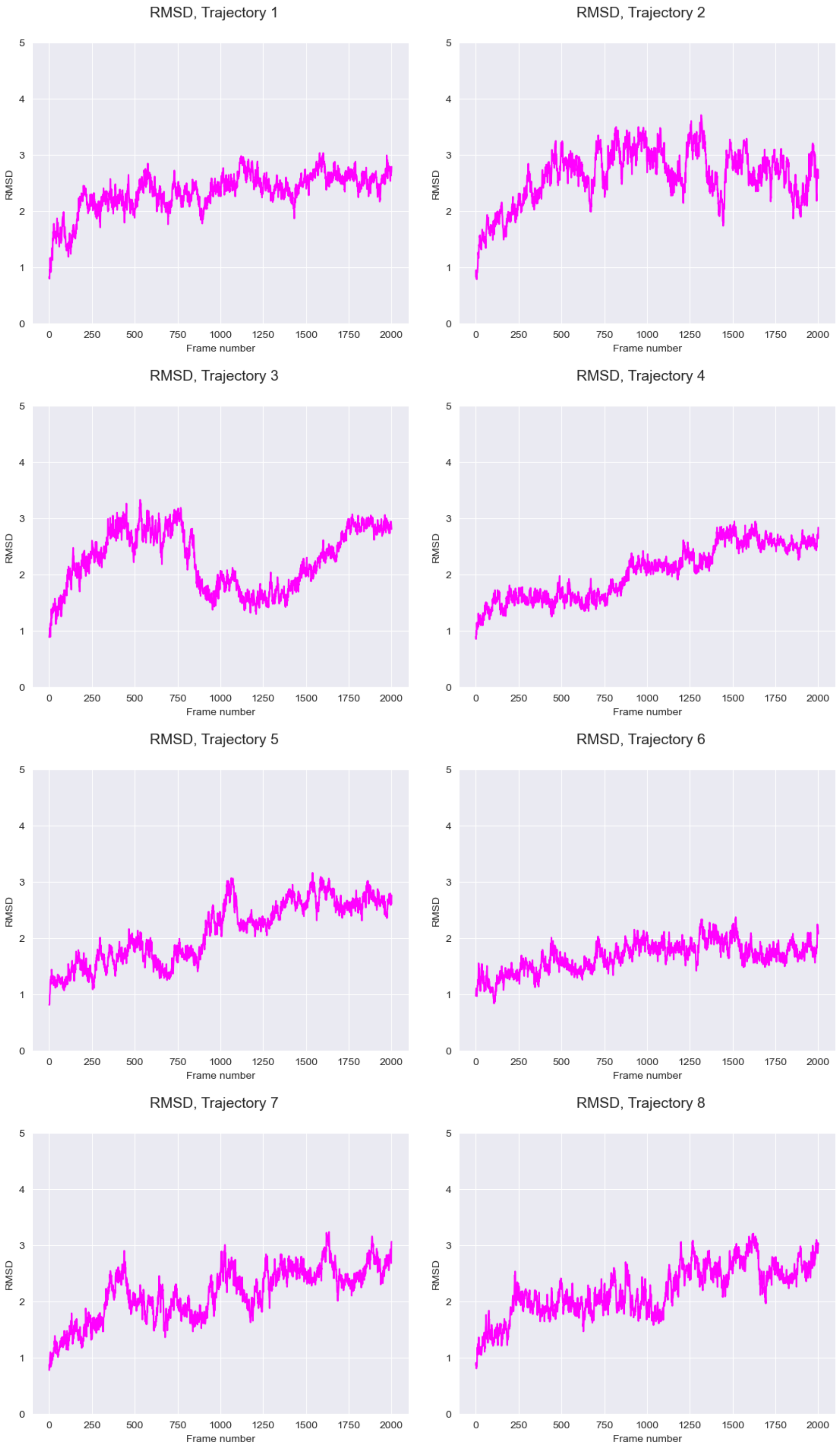
RMSD, Trajectory 1
RMSD, Trajectory 2
RMSD, Trajectory 3
RMSD, Trajectory 4
RMSD, Trajectory 5
RMSD, Trajectory 6
RMSD, Trajectory 7
RMSD, Trajectory 8
RMSD
Frame number
0
1
2
3
4
5
250
500
750
1000
1250
1500
1750
2000

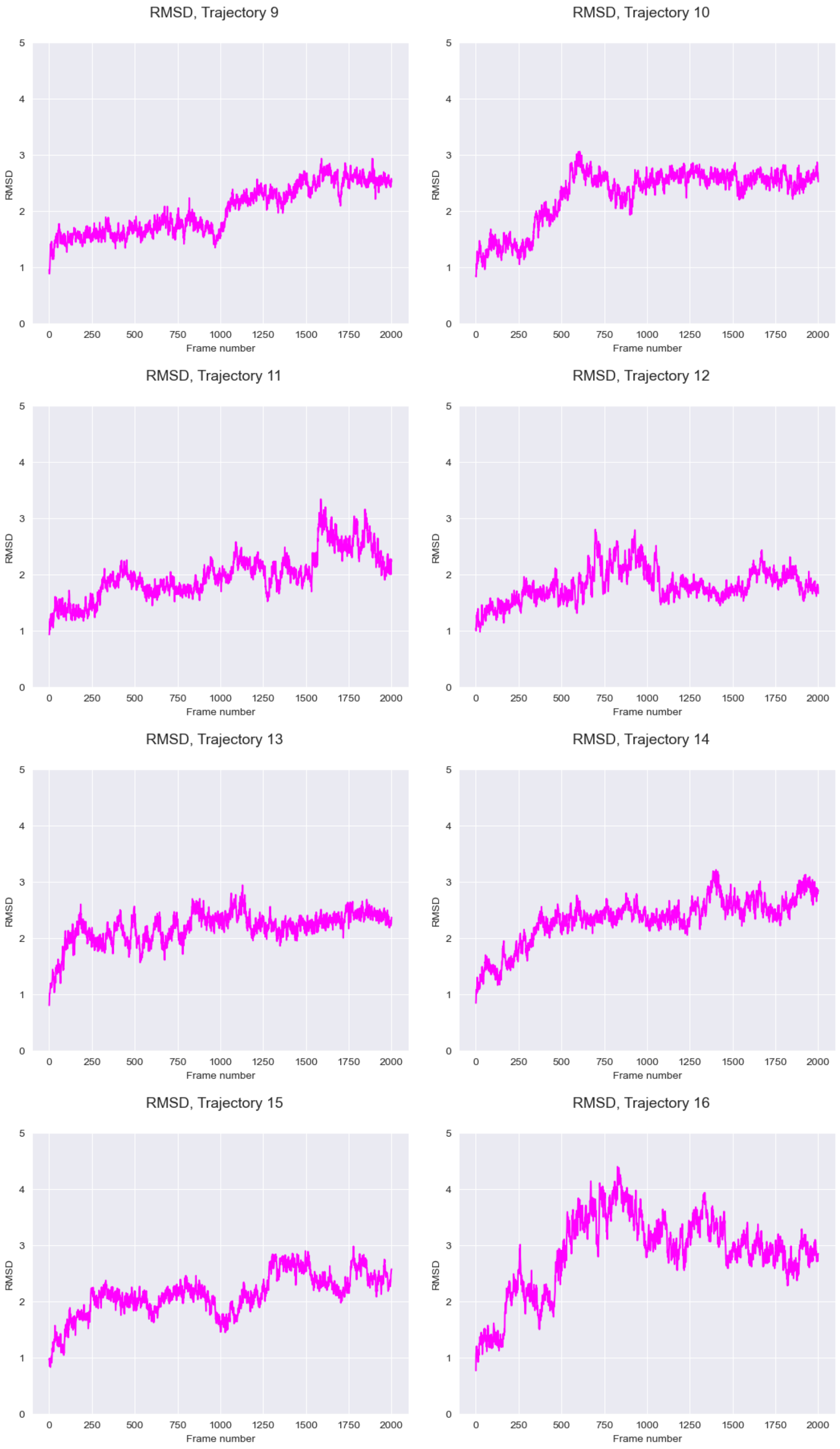
RMSD, Trajectory 9
RMSD, Trajectory 10
RMSD, Trajectory 11
RMSD, Trajectory 12
RMSD, Trajectory 13
RMSD, Trajectory 14
RMSD, Trajectory 15
RMSD, Trajectory 16
RMSD
Frame number
0
1
2
3
4
5
0
250
500
750
1000
1250
1500
1750
2000

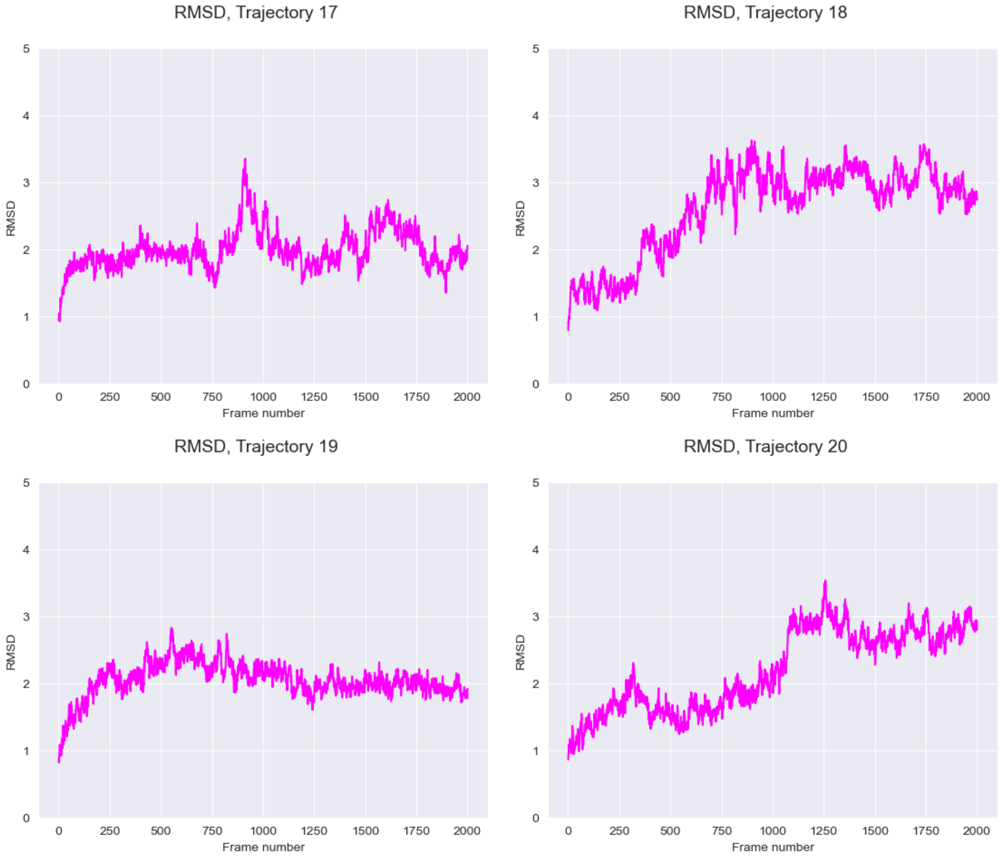


Figure S5: *c-kit1* GQ RMSD for each of the 20 independent MD simulations starting from an empty ionic channel.

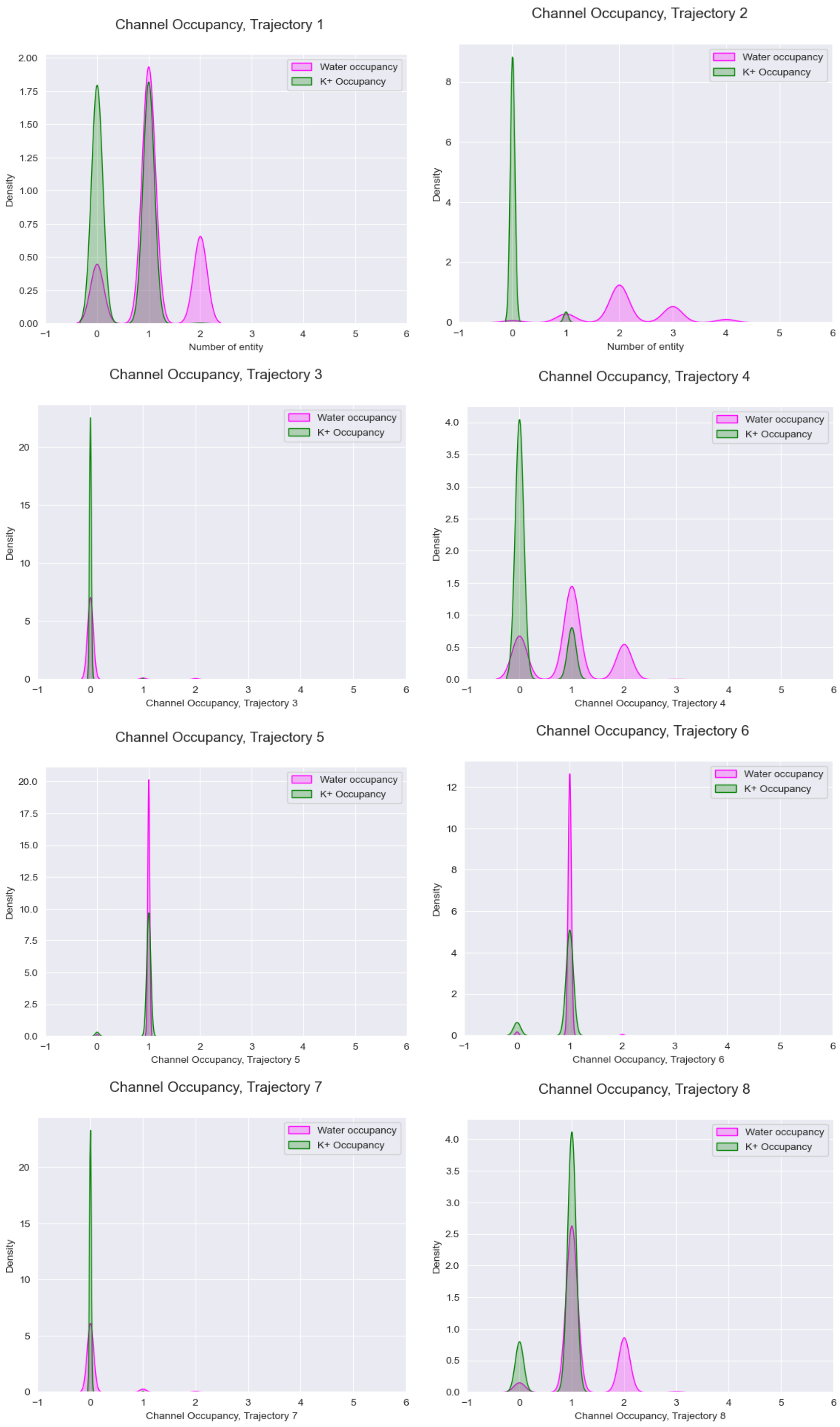
Channel Occupancy, Trajectory 1
Water occupancy
K+ Occupancy
Density
Number of entity
Channel Occupancy, Trajectory 2
Water occupancy
K+ Occupancy
Density
Number of entity
Channel Occupancy, Trajectory 3
Water occupancy
K+ Occupancy
Density
Channel Occupancy, Trajectory 3
Channel Occupancy, Trajectory 4
Water occupancy
K+ Occupancy
Density
Channel Occupancy, Trajectory 4
Channel Occupancy, Trajectory 5
Water occupancy
K+ Occupancy
Density
Channel Occupancy, Trajectory 5
Channel Occupancy, Trajectory 6
Water occupancy
K+ Occupancy
Density
Channel Occupancy, Trajectory 6
Channel Occupancy, Trajectory 7
Water occupancy
K+ Occupancy
Density
Channel Occupancy, Trajectory 7
Channel Occupancy, Trajectory 8
Water occupancy
K+ Occupancy
Density
Channel Occupancy, Trajectory 8

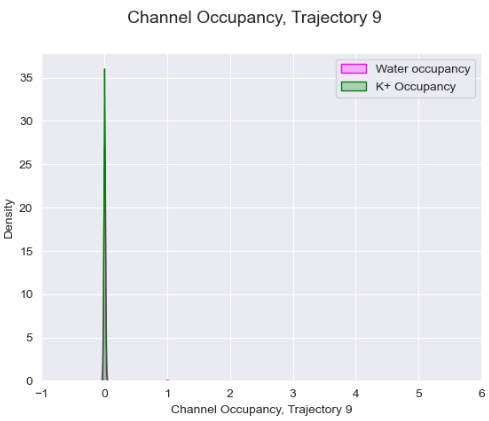
Channel Occupancy, Trajectory 9
Water occupancy
K+ Occupancy
Density
0
5
10
15
20
25
30
35
−1
0
1
2
3
4
5
6
Channel Occupancy, Trajectory 9

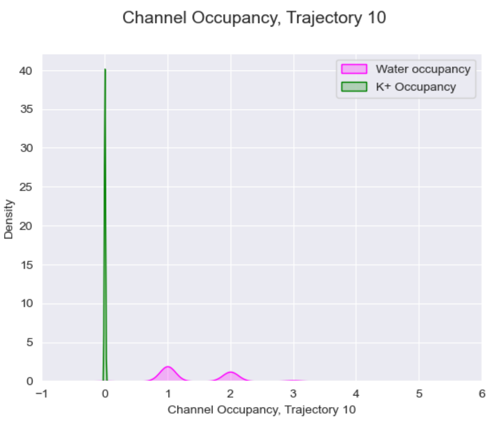
Channel Occupancy, Trajectory 10
Water occupancy
K+ Occupancy
Density
0
5
10
15
20
25
30
35
40
−1
0
1
2
3
4
5
6
Channel Occupancy, Trajectory 10

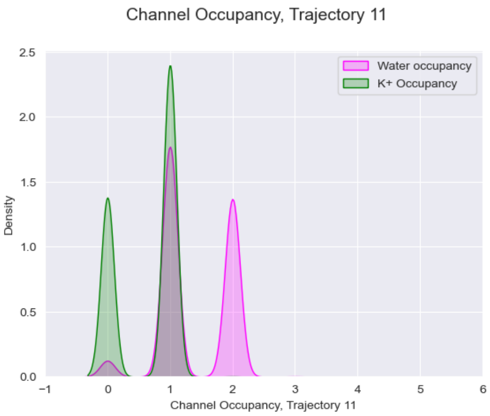
Channel Occupancy, Trajectory 11
Water occupancy
K+ Occupancy
Density
0.0
0.5
1.0
1.5
2.0
2.5
−1
0
1
2
3
4
5
6
Channel Occupancy, Trajectory 11

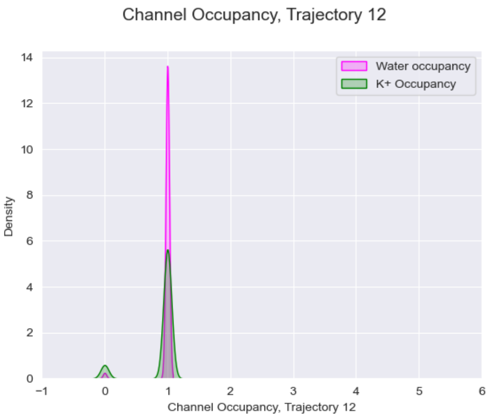
Channel Occupancy, Trajectory 12
Water occupancy
K+ Occupancy
Density
0
2
4
6
8
10
12
14
−1
0
1
2
3
4
5
6
Channel Occupancy, Trajectory 12

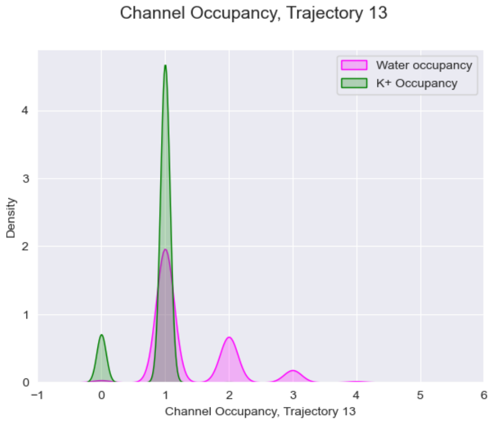
Channel Occupancy, Trajectory 13
Water occupancy
K+ Occupancy
Density
0
1
2
3
4
−1
0
1
2
3
4
5
6
Channel Occupancy, Trajectory 13

l

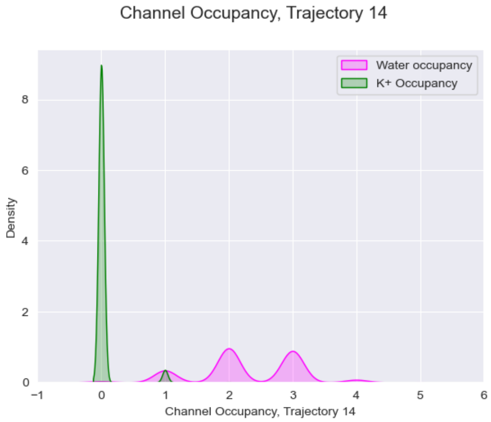
Channel Occupancy, Trajectory 14
Water occupancy
K+ Occupancy
Density
0
2
4
6
8
−1
0
1
2
3
4
5
6
Channel Occupancy, Trajectory 14

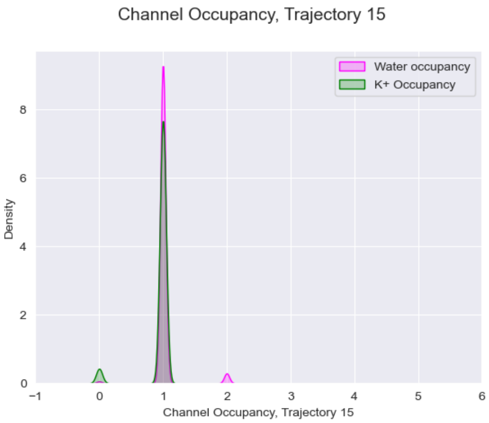
Channel Occupancy, Trajectory 15
Water occupancy
K+ Occupancy
Density
0
2
4
6
8
−1
0
1
2
3
4
5
6
Channel Occupancy, Trajectory 15

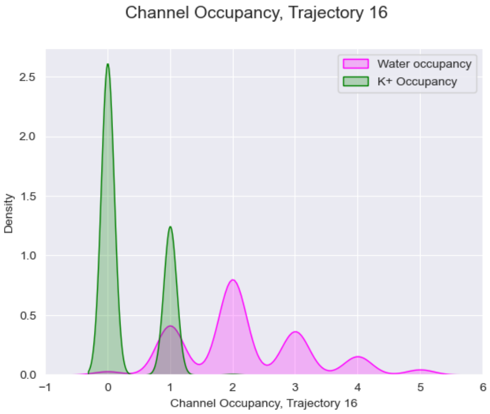
Channel Occupancy, Trajectory 16
Water occupancy
K+ Occupancy
Density
0.0
0.5
1.0
1.5
2.0
2.5
−1
0
1
2
3
4
5
6
Channel Occupancy, Trajectory 16

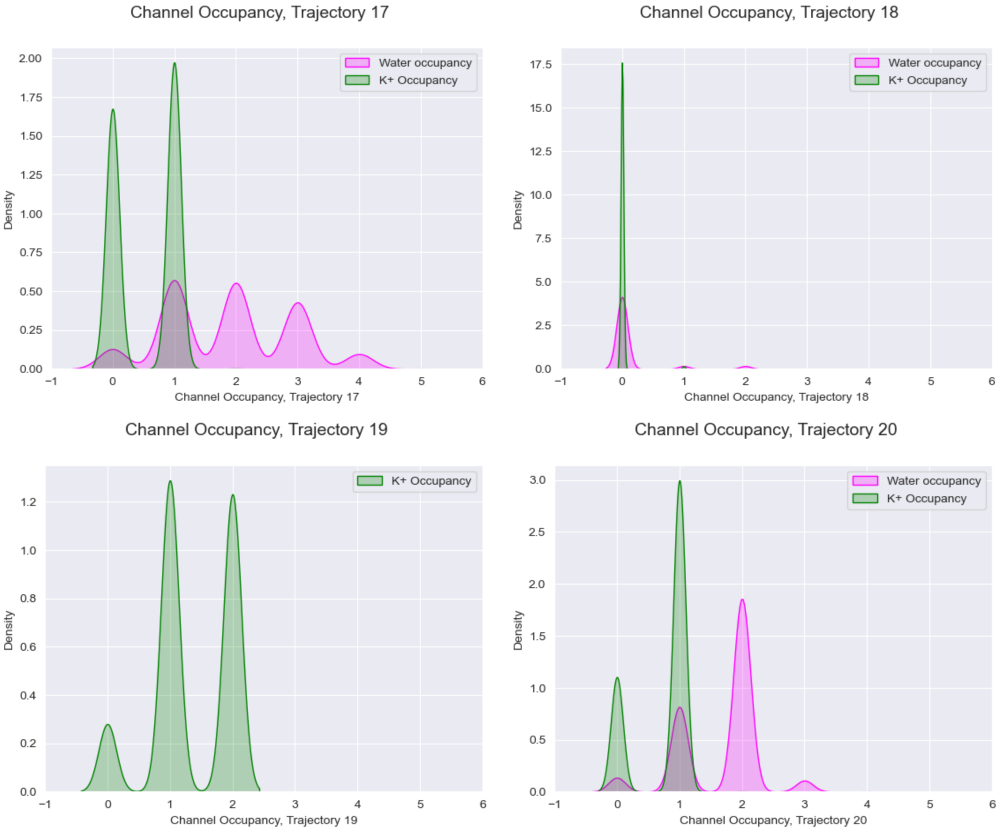


Figure S6: Probability density function representation of *ckit1* GQ channel occupancy with water molecules, presented in green and with $K^+$ cations, presented in fushia, thoughout 20 independent MD trajectories starting from an empty ionic channel.

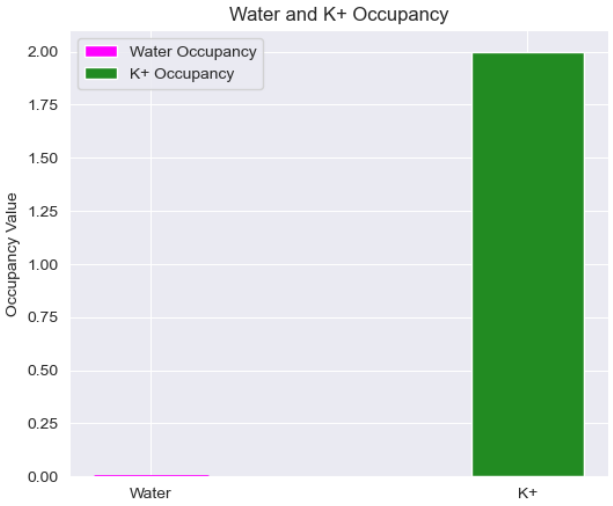


Figure S7: *ckit1* GQ channel occupancy. Water molecules, presented in green and with $K^+$ cations, presented in fushia, thoughout 800ns CMD trajectory.

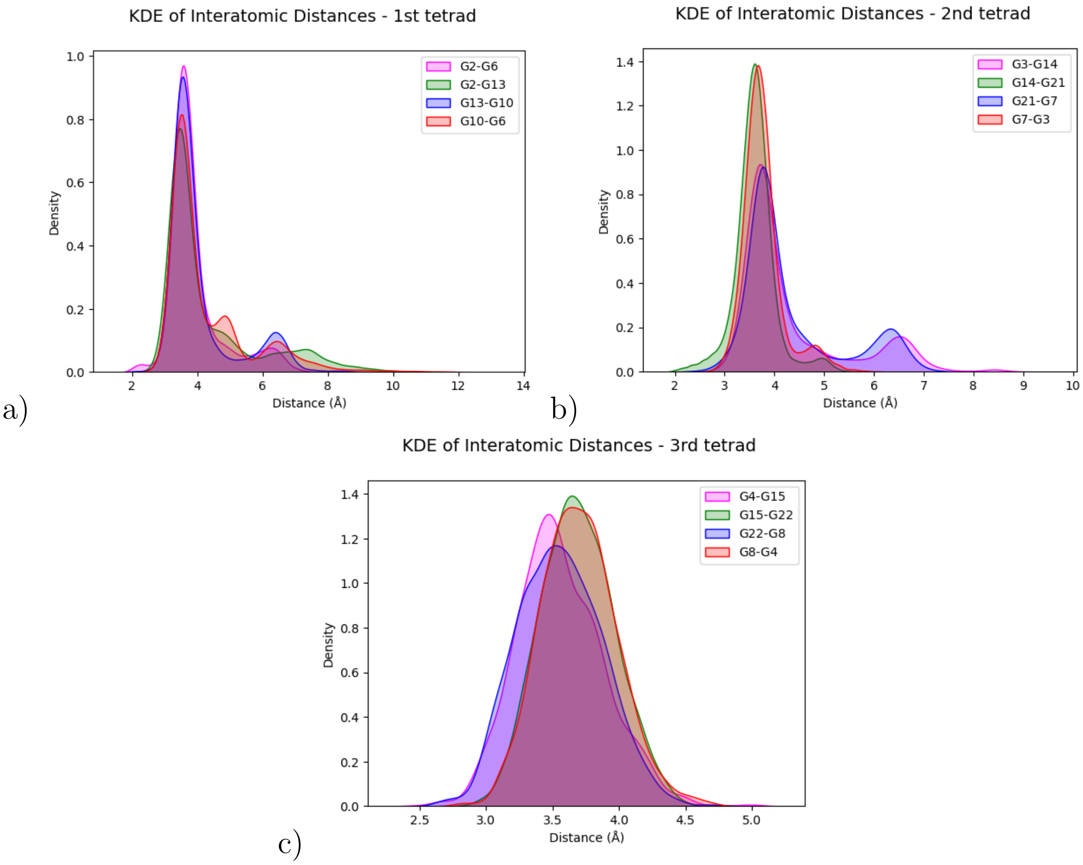


Figure S8: Probability density function representation of the distances between N7 and H1 of 2 adjacent guanines within a same tetrad.

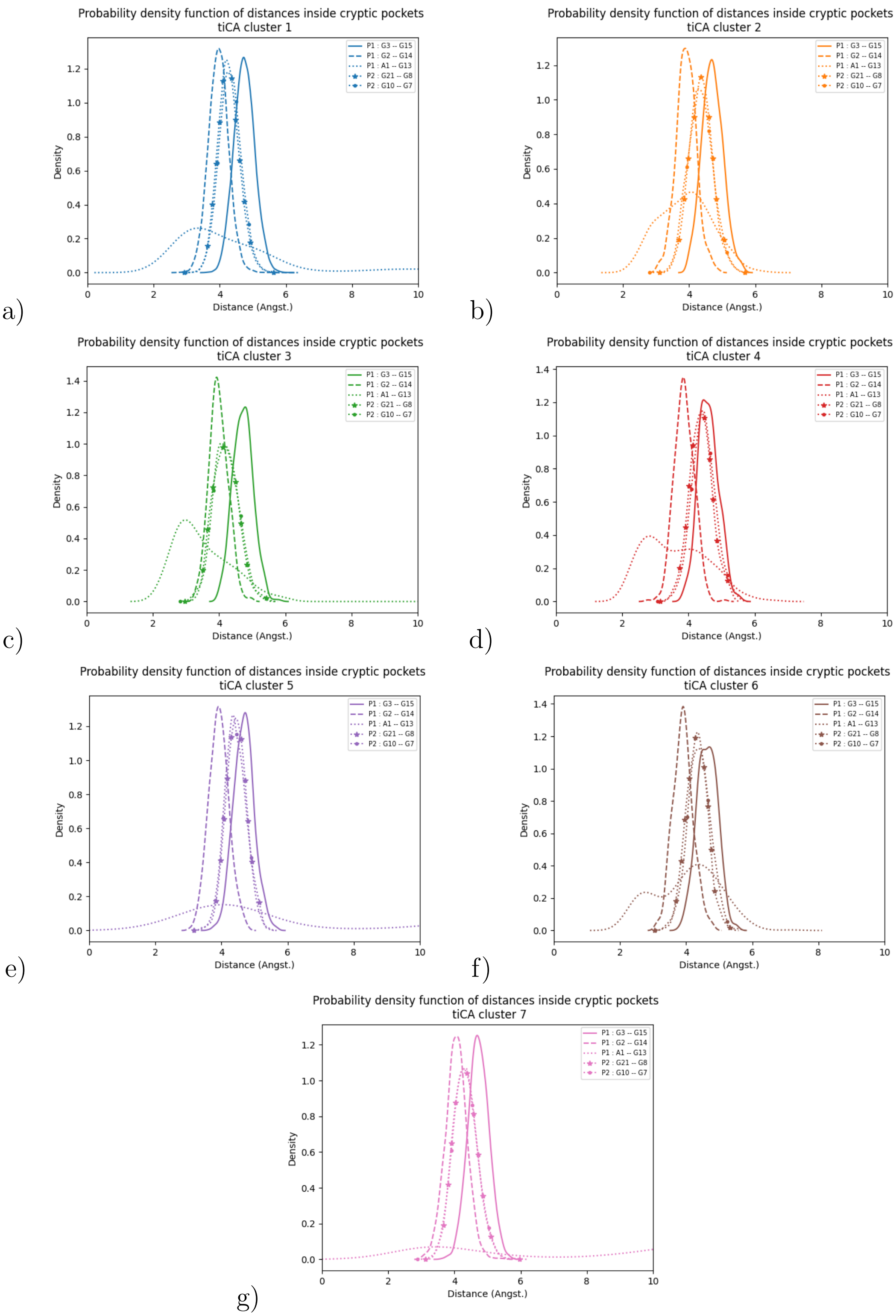


Figure S9: Probability density function representation of the distances between nucleotides inside the cryptic pockets P1 and P2 defined in *c-kit 1* GQ . See text for definition

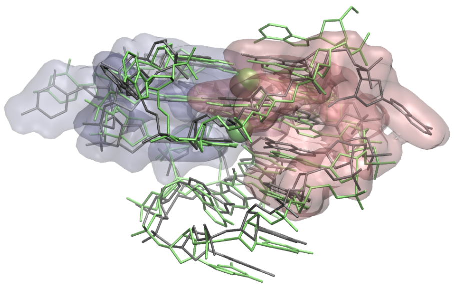

a)

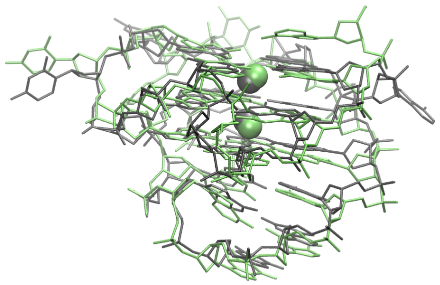

b)

Figure S10: Representation of superposed *c-kit 1*
GQ 3D structures and the corresponding cations inside the GQ channel. We represent in gray the initial conformation identical to the X-Ray, and in green the most representative cluster conformation. We show in a) the identified pockets P1 and P2 in pink and iceblue surface respectively. The same represention is shown in b) without the pocket surface for a better visibility of the ions inside the channel.

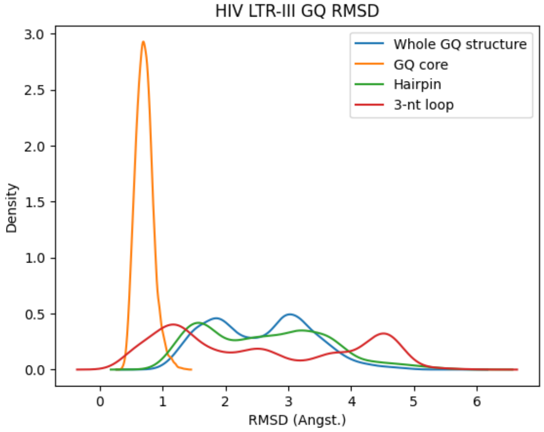


a)

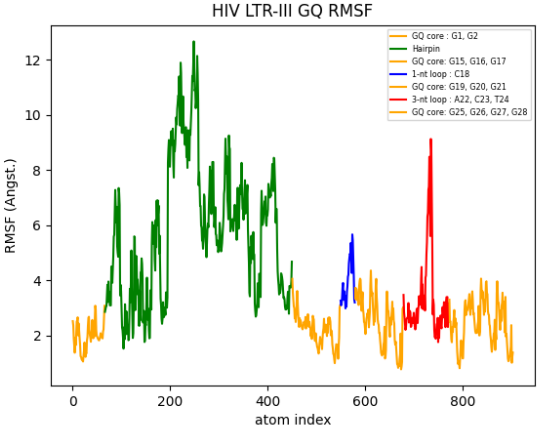


b)

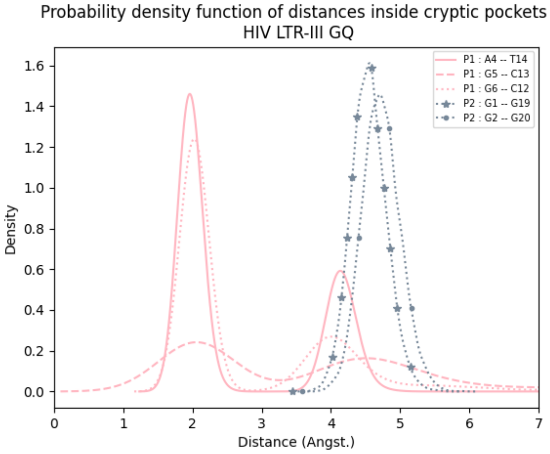


c)

Figure S11: a) RMSD and b) RMSF of HIV LTR-III GQ. c) Probability density function of A1–T14, G5–C13, G6–C12 distances inside P1 and of G2–G19 and G1–G20 inside P2.

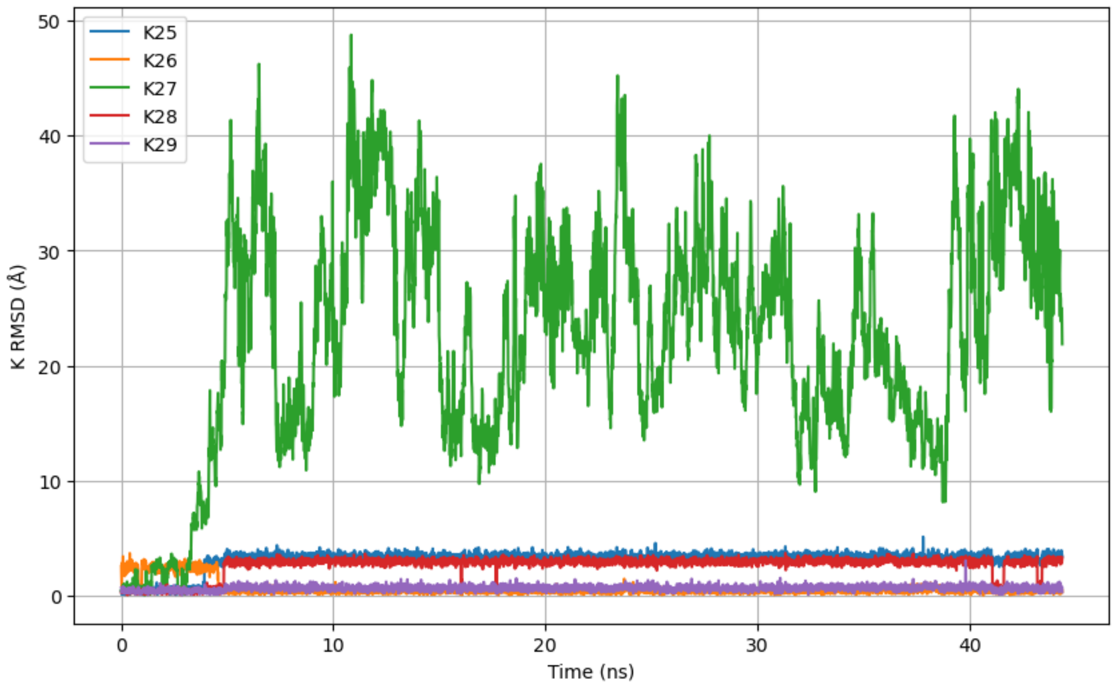


a)

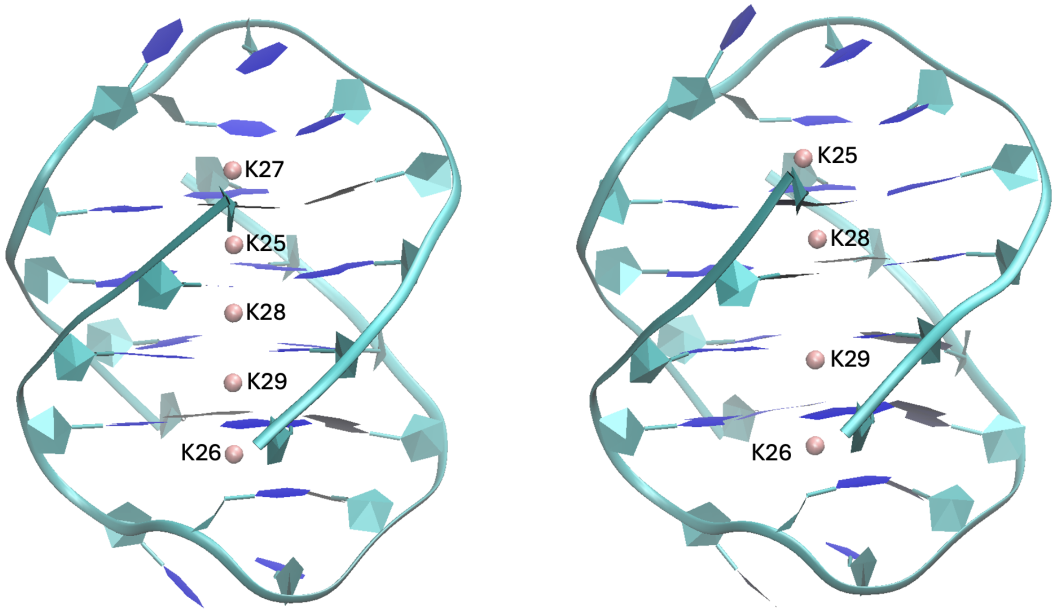


b)

Figure S12: a) The distance from the original position ofeach of the $K^+$ ions in the GQ channel of 1JRN over 50 ns and b)$K^+$ ion positioning in the GQ channel of 1JRN at the beginning (left) and end (right) of the 50 ns MD simulation.
